# ganon: precise metagenomics classification against large and up-to-date sets of reference sequences

**DOI:** 10.1101/406017

**Authors:** Vitor C. Piro, Temesgen H. Dadi, Enrico Seiler, Knut Reinert, Bernhard Y. Renard

## Abstract

**Motivation:** The exponential growth of assembled genome sequences greatly benefits metagenomics studies. However, currently available methods struggle to manage the increasing amount of sequences and their frequent updates. Indexing the current RefSeq can take days and hundreds of GB of memory on large servers. Few methods address these issues thus far, and even though many can theoretically handle large amounts of references, time/memory requirements are prohibitive in practice. As a result, many studies that require sequence classification use often outdated and almost never truly up-to-date indices.

**Results:** Motivated by those limitations we created ganon, a k-mer based read classification tool that uses Interleaved Bloom Filters in conjunction with a taxonomic clustering and a k-mer counting/filtering scheme. Ganon provides an efficient method for indexing references, keeping them updated. It requires less than 55 minutes to index the complete RefSeq of bacteria, archaea, fungi and viruses. The tool can further keep these indices up-to-date in a fraction of the time necessary to create them. Ganon makes it possible to query against very large reference sets and therefore it classifies significantly more reads and identifies more species than similar methods. When classifying a high-complexity CAMI challenge dataset against complete genomes from RefSeq, ganon shows strongly increased precision with equal or better sensitivity compared with state-of-the-art tools. With the same dataset against the complete RefSeq, ganon improved the F1-Score by 65% at the genus level. It supports taxonomy- and assembly-level classification, multiple indices and hierarchical classification.

**Availability:** The software is open-source and available at: https://gitlab.com/rki_bioinformatics/ganon

## 1 Introduction

Reference- and taxonomy-based short read classification is a fundamental task in metagenomics. Defining the origin of each read from an environmental sample, which can be done during (Tausch *et al.*, 2018) or after sequencing, is usually the first step prior to abundance estimation, profiling and assembly. Over the last years many tools have been specifically developed for this task (Oulas *et al.*, 2015; McIntyre *et al.*, 2017; Lindgreen *et al.*, 2016; Peabody *et al.*, 2015; Sczyrba *et al.*, 2017) with different strategies to achieve good performance classifying a large amount of short reads against a predefined and static set of reference sequences. Many of those approaches are taxonomy-based (Balvočiūtė and Huson, 2017) and use this classification to better understand the composition of samples.

The amount of complete or draft genomic sequences in public repositories is rapidly growing due to advances in genome sequencing, improvements in read quality, length and coverage and also better algorithms for genome assembly. In addition, many partial and complete genome sequences come directly from metagenome-assembled genomes (Parks *et al.*, 2017; Mukherjee *et al.*, 2017; Tully *et al.*, 2018), a technique that boosts the growth of public repositories. This considerable amount of references poses a sizeable challenge for current tools that, in general, are not designed to deal with such amounts of data (Nasko *et al.*, 2018). They also increase the already high computational cost of assigning millions of short reads to taxonomic targets.

Using GenBank (Benson *et al.*, 2018) and RefSeq (Haft *et al.*, 2018) repositories as an example, we see an exponential data growth (Supplementary Material 1 - Figure 1). Within an interval of two and a half years (from June 2015 to December 2018) the RefSeq Microbial of Complete Genomes (CG) grew more than four times, with 2.5 times more species represented in the most recent set (1529 to 3850). Looking at the same data point (end of 2018), the complete RefSeq Microbial has >12 times base pairs and >5 times species compared to the CG set. These data exemplify that databases are growing fast and the variation among them is significant. These repositories are becoming too big to be analyzed by standard hardware and if the observed growth continues, all this wealth of data will be constrained to just a few groups with resources available to process them.

The choice of the data to perform reference-based classification is an important step and a known issue in metagenomics (Breitwieser *et al.*, 2017). As a rule of thumb, the more sequences the better the classification. But even complete sets of sequences are not evenly distributed throughout the taxonomic tree, such that different taxa are represented in different levels of quantity and quality. In addition, most of the taxa are still unknown and do not have any genomic sequence or entry in the taxonomic tree. This requires the tools to consistently remain up to date with the latest releases of public repositories, a task that is not trivial when dealing with very large amounts of sequences. Most of the tools lack the ability to update their own indices and databases, and currently many analyses are performed with outdated resources.

For example, the RefSeq Microbial repository from the beginning of 2018 is 10% less taxonomic diverse than it is today (mid. 2019). An even older RefSeq release from June 2015 lacks 27% of today’s taxonomic diversity. Further, a commonly used subset of RefSeq, the microbial complete genomes, covers only 15% of the available diversity of the full repository (December 2018). As an example, the latest release of kraken’s (Wood and Salzberg, 2014) MiniKraken database (as of 18-Oct-2017) based on complete bacterial, archaeal, and viral genomes, although helpful to obtain fast insights on community composition, comprises only 11% of the total taxonomic diversity available on the latest RefSeq release from January 4th 2019. Metagenomics analyses based on those releases are prone to underperform and miss potential species of interest. However, the use of outdated references or “pre-built” indices is still common practice (Li *et al.*, 2018). Most methods are able to build custom databases but unable to update them. Weekly or daily updates with the most recent data are almost impossible given the time requirements to re-build those indices.

The sequence classifiers MetaPhlAn (Truong *et al.*, 2015) and Kaiju (Menzel *et al.*, 2016) created alternatives to cover most of the diversity contained in public sequence repositories by selecting a subset of marker genes and protein sequences, respectively. On one hand, those methods are very powerful, such that they provide fast and precise community profiles given their reduced index sizes. On the other hand, when analyzing whole genome sequences of complex environments, organisms with low abundance are easily missed due to their lack of complete genomic coverage. In addition, current methods using complete genome sequences struggle with the present amount of available data (Nasko *et al.*, 2018).

Given these limitations, we developed ganon, a new reference and taxonomy-based short read classification tool for metagenomics. Ganon uses Interleaved Bloom Filters (IBF) (Dadi *et al.*, 2018) to represent very large amounts of sequences into a searchable index. This enables the indexing of large sets of references (e.g. complete RefSeq) in faster time and with low memory consumption, consequently improving read classification for whole metagenomics sequencing experiments. Ganon also provides updatable indices, which can incorporate new released sequences in short time. The classification method, which is based on the k-mer counting lemmaandaprogressive filtering step, improves sensitivity and precision compared to state-of-the-art tools when using larger sets of references. Ganon was developed in C++ using the SeqAn library (Reinert *et al.*, 2017) and Python. The code is open source and freely available from: https://gitlab.com/rki_bioinformatics/ganon

## 2 Methods

### 2.1 Overview

Ganon classifies reads against a set of reference sequences to find their exact or closest taxonomic origin. The method can also work in a further specialized level (e.g. assembly). Clustering and indexing steps are necessary before classification, where the reference sequences will be grouped into taxonomic groups and processed into a searchable index. Ganon indices store all k-mers present in the reference sequences into a specialized type of Bloom filter. Once the index is created, ganon classifies the reads based on the k-mer counting lemma together with a post-filtering step providing a unique or multiple classifications for each read. Multiple classifications are solved optionally with the lowest common ancestor (LCA) algorithm (Huson *et al.*, 2007). The following sections will further explain each of these steps in detail.

### 2.2 Indexing

Ganon indices are based on the k-mer content of the reference sequences, in other words, it uses all possible substrings of length *k* of the given sequences. Instead of using standard methods for k-mer storage, which can have high memory and space consumption when *k* is high (> 15), we opted for Bloom filters (Bloom, 1970), a space-efficient probabilistic data structure. Since the goal of the tool is to classify sequences based on their taxonomic origin, multiple Bloom filters would be necessary to represent each distinct group of sequences belonging to a certain taxonomic level (e.g. species). This approach provides a straightforward but impractical solution since it requires classification against multiple filters. This is solved by interleaving the Bloom filters, a technique previously described for the DREAM-Yara tool (Dadi *et al.*, 2018) and also part of the SeqAn library (Reinert *et al.*, 2017). TaxSBP is used to separate sequences into taxonomic groups and to distribute them better into equal-sized clusters.

#### 2.2.1 TaxSBP

TaxSBP [https://github.com/pirovc/taxsbp] uses the NCBI Taxonomy database (Federhen, 2012) to generate clusters of sequences that are close together in the taxonomic tree. It does this based on an implementation of the approximation algorithm for the hierarchically structured bin packing problem (Codenotti *et al.*, 2004). As defined by Codenotti *et al.* this clustering method “[…] can be defined as the problem of distributing hierarchically structured objects into different repositories in a way that the access to subsets of related objects involves as few repositories as possible”, where the objects are sequences assigned to taxonomic nodes of the taxonomic tree. Sequences are clustered together into groups limited by a maximum sequence length size of its components. Splitting sequences into smaller chunks with overlapping ends is supported. TaxSBP supports one level of specialization after the leaf nodes of the tree, making it possible to further cluster sequences by strain or assembly information that is not directly contained in the NCBI Taxonomy database (Figure 1 A). TaxSBP can also pre-cluster members of a certain taxonomic level, preventing them to be split among clusters. It can further generate clusters with exclusive ranks, which are guaranteed to be unique in their cluster. The tool was developed alongside the distributed indices concept (Dadi *et al.*, 2018) and supports the update of pre-generated clusters. Since TaxSBP uses the “pre-clustered” taxonomic tree information, the algorithm is very efficient and requires very few computational resources, thus having potential use in many other bioinformatics applications.

**Fig. 1.**
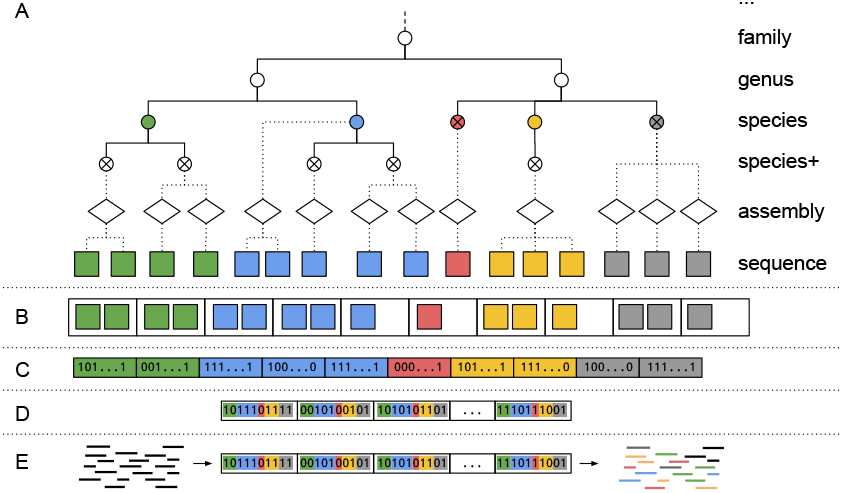
Ganon methodology overview. A) Empty circles are inner nodes of the tree; “x” circles are leaf nodes (referenced in this manuscript as taxid nodes); full lines represent taxonomic relations, dotted lines represent the extension of the taxonomy to the assembly and sequence levels. Species+ represents all taxonomic groups that are more specific than species with species in the lineage (e.g. subspecies, species group, no rank). B) A toy example of sequences clustered by species into equal-sized groups, performed by TaxSBP C) Sequences are fragmented into k-mers and with a given number of hash functions, those k-mers are inserted into equal-sized bit-vectors (Bloom Filters) D) The Interleaved Bloom Filter, representing the previously generated bit-vectors with each bit interleaved. E) Classification of short reads (black lines) against the IBF. Reads are fragmented into k-mers, counted with the same hash functions against the IBF, filtered and assigned to one or more species followed by LCA assignment for multiple matches.

#### 2.2.2 IBF

A Bloom filter is a probabilistic data structure that comprises a bit vector and a set of hash functions. Each of the functions maps a key value (k-mer in our application) to one of the bit positions in the vector. Collisions in the vector are possible, meaning that distinct k-mers can be set to the same bit positions in the vector. Those overlaps can be avoided with a larger bit vector, thus reducing the probability of false positives.

An Interleaved Bloom Filter (IBF) is a combination of several (*b*) Bloom filters of the same size (*n*) with the same hash functions into one bit vector (Figure 1 D). Each *i*-th bit of every Bloom filter is interleaved, resulting in a final IBF of size *b* * *n*. Querying in this data structure is possible by retrieving the sub-bit vectors for every hash function and merging them with a logical *AND* operation, which will result in a final bit vector indicating the membership for the query, as depicted in Figure 2 in the DREAM-Yara manuscript by (Dadi *et al.*, 2018).

**Fig. 2.**
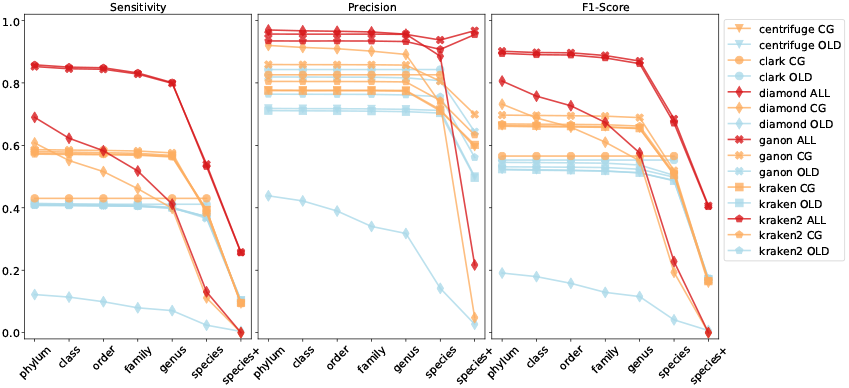
Cumulative-based precision, sensitivity and F1-Score values at all ranks for the simulated reads against all evaluated reference sets (blue = RefSeq-OLD, orange = RefSeq-CG-top-3, red = RefSeq-ALL-top-3)

Aiming at the classification based on taxonomic levels (e.g. species, genus,…) or assembly level, TaxSBP is set to cluster the input sequences into exclusive groups (Figure 1 B). Every group will contain only sequences belonging to the same taxon or assembly unit, but the same unit can be split into several groups. Groups are limited by a predefined threshold of the sum of the base pair length of its elements and sequences can be sliced into smaller pieces to better generate equal sized clusters.

Each of those clusters will correspond to a single Bloom filter that is interleaved in a final IBF (Figure 1 C-D). Here a trade-off between the number of groups, their maximum size and the k-mer content of each group is important. The false positive rate of a Bloom filter depends mainly on its bit vector size and the number of inserted elements. In general, the more base pairs a particular cluster has, the higher the number of distinct k-mers. This requires the Bloom filter to be bigger in order to achieve low false positive rates when querying. In ganon indices, the group with the most unique k-mers will define the size and the maximum false positive rate of the final IBF since all groups have to be equal-sized by definition. Thus the previous clustering step is crucial to achieve a good trade-off between the number of groups, their sizes and k-mer content. The lower the taxonomic level, the more fragmented the clusters. For example: if a reference set has 2000 species groups, there will be at least the same number of clusters when building at the species level. The higher the taxonomic level, the fewer the number of clusters, since they can be grouped together, thereby producing smaller filters. This trade-off and parameterization is automatically calculated by ganon, with a single option to define the maximum memory available to build an index.

The IBF has an inherent capability of updating since it is fragmented into many sub-parts. Adding new sequences to a previously generated IBF is as easy as setting the bit positions of the k-mers from the new sequences to their known clusters or appending new clusters to the existing filter. To remove sequences from the IBF, all bit positions of the updated cluster are set to zero and the cluster is re-created from the updated content.

The IBF is the main data structure for ganon indices to perform alignment-free classification while DREAM-Yara, the tool that originally proposed the IBF, is a read mapper that uses the same data structure to filter reads to further perform distributed alignment. At the end of the building process, the ganon index will consist of an IBF based on a maximum classification level chosen (taxonomic rank or assembly) and auxiliary files for the classification step.

### 2.3 Classifying

The read classification is based on the well-studied k-mer counting lemma (*q*-gram lemma (Jokinen and Ukkonen, 1991; Reinert *et al.*, 2015)). All k-mers from given reads are looked up on the indices previously generated. If a minimum number of matches between the read and the reference is achieved, a read is considered classified. Based on incremental error rates, multiple classifications for each read are filtered out and only the best ones are selected. When the filtering cannot define a single origin for a read, an optional LCA step is applied to join multiple matching reads into their lowest common ancestor node in the taxonomic tree.

#### 2.3.1 K-mer counting lemma

The k-mer counting lemma can be defined as the minimum number of k-mer sequences of a read that should match against reference k-mers in order to be considered present in a set with a certain number of errors allowed. Given a read, *R*, with length *l*, the number of possible k-mers with length *k* in this read can be defined as:

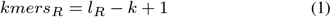

Based on the *q*-gram lemma, an approximate occurrence of *R* in a set of references has to share at least

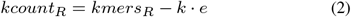

k-mers, where e is the maximum number of errors/mismatches allowed.

#### 2.3.2 Filtering

A read can be assigned to multiple references with different error rates, thus a filtering step is necessary to decrease the number of false assignments. The applied k-mer counting lemma provides k-mer counts for each read against the reference sequences. From this count it is possible to estimate the number of mismatches a read has. For example: for *k* =19 and *length* = 100, a read with 50 19-mers matching a certain reference will optimally have 2 mismatches. This calculation can be achieved by solving the Equation 2 equation for *e*.

Assuming that reads with fewer mismatches have a higher chance of being correct, the following filtering is applied: first, only matches against references with no mismatches are kept (all k-mers matching). If there are no such assignments, matches with only 1 error are kept. If there are none, matches with only 2 errors are kept and so on up to the maximum number of errors allowed (*e* in Equation 2). Similar filtration methods (also known as mapping by strata) were previously used in read mappers such as Yara (Dadi *et al.*, 2018). If a read is classified in several references within the same range of errors, they are all reported since it is not possible to define which one has a higher chance of being correct based on the k-mer count information. Given our clustering approach, some groups can share the same identification target (e.g. one species was split in two or more clusters due to a large amount of sequences). These cases are treated specially by reporting only the match with more k-mer similarities since they belong to the same classification group.

Ganon also provides a way to further filter the unique classifications with a different error rate for reads that matched against just one reference group. This filter will be applied after the standard filtration and will reclassify a low scored read to its parent taxonomic level if it scores below a certain threshold. This can be applied for filtering at low levels (e.g. assembly) since the classification in those levels should be more precise with less mismatches. This feature is also useful to avoid classifications that only happen due to a lack of related genomes (e.g. a low score match on the only representative species of a lineage).

In summary, ganon indices represent groups of reference sequences clustered by taxonomy or assembly group. All k-mers from the reads are extracted and compared against an index by applying the k-mer counting lemma to select candidates. This is done based on a user defined optimal number of errors. All matches within the error rate are filtered and one or more matches are reported. At the end, an optional LCA method can be applied for reads with multiple matches with a more conservative and less precise taxonomic classification, thus resulting in one match for each read. Additionally, ganon supports classification based on multiple indices in a user-defined hierarchy, with independent error rates for each index (Supplementary Material 1 - Section 3.4).

## 3 Results

We evaluated ganon against a set of well-established methods from recent benchmarks (McIntyre *et al.*, 2017; Lindgreen *et al.*, 2016; Sczyrba *et al.*, 2017) that performs short read classification and supports indexing of large sets of reference sequences. The aim here is to compare in equal conditions the methods regarding input data, reference sequences and taxonomy. We compared ganon against kraken (Wood and Salzberg, 2014), one of the most used k-mer based methods for metagenomics short read classification and its newer version, kraken2 (Wood *et al.*, 2019). We also included krakenuniq (Breitwieser *et al.*, 2018), which uses the basic kraken algorithm and also allows classification on more specific levels after taxonomic assignments (e.g. up to assembly or sequence level). We further compare the results against centrifuge (Kim *et al.*, 2016) that uses the Burrows-Wheeler transform (BWT) and the Full-text index in Minute space (FM-)index for indexing and aims to reduce the index size by compressing similar sequences together. Clark (Ounit *et al.*, 2015), another k-mer approach that uses common k-mers between reference sequences was also evaluated. Diamond (Buchfink *et al.*, 2015) an alignment tool for short DNA sequencing reads against protein reference databases was also included. Here we consider only the direct read classification capabilities of the tools. Further functionalities such as the estimation of a presence of a certain organism or abundance estimation were not covered. All steps performed in the evaluation were compiled in a benchmark pipeline (version 1.1.0) available from https://github.com/pirovc/ganon_benchmark.

Ganon and the other evaluated tools are reference-based, meaning all classifications are made based on previously generated sequence data. The choice of the underlying database is therefore crucial. We use the same sequences and taxonomic database version for all tools when creating their indices to guarantee a fair comparison. The NCBI RefSeq repository was the chosen source of sequences since it provides a curated and very extensive set of references. Two subsets of RefSeq were extracted: a set of only complete genomes from the groups Archaea, Bacteria, Fungi and Viral (RefSeq-CG) and a complete set of all genomes from the same groups (RefSeq-ALL) both dating from 19-December-2018 (Table 1). Genomic DNA data was obtained for all tools. Protein sequence data from annotated genome assemblies was obtained for diamond (Supplementary Material 1 - Table 5). Taxonomic information was obtained at the same dates as the sequences. Additionally, an old set of only Bacterial complete genomes from 02-June-2015 (RefSeq-OLD) was included to evaluate the tool’s performance on an outdated and less diverse set of references.

**Table 1.** Genomic DNA of reference sequences used for evaluations. Protein data information can be found in the Supplementary Material 1 - Table 5. Detailed information of each dataset can be found in the Supplementary Material 1 - Section 2.5.1. Data was downloaded using https://github.com/pirovc/genome_updater

|  | Base pairs | # assemblies | # sequences |
| --- | --- | --- | --- |
| RefSeq-OLD | 9,632,441,987 | 3,042 | 5,242 |
| RefSeq-CG | 46,986,899,184 | 19,623 | 33,029 |
| RefSeq-ALL | 587,607,072,429 | 147,713 | 15,201,684 |

The selected reference sets contain over-represented taxonomic groups with several assemblies for a single species. For example: the *Escherichia coli* species group is represented by 634 assemblies, accounting for almost 7% of all base pairs in the RefSeq-CG. This is even more pronounced on RefSeq-ALL, with 13,259 *E. Coli* assemblies representing more than 11% of the base pairs in the whole set. In RefSeq-CG, the 92 most over-represented species have as many base pairs as the remaining 11,372 species. In RefSeq-ALL this ratio is 14 to 29,047 (Supplementary Material 1 - Figure 2). This unbalanced distribution of references may not only bias analysis but also introduces redundancy to the set when aiming classification at taxonomic levels. Therefore, when not classifying at assembly level, we removed over-represented assemblies from our reference set, keeping only the 3 biggest assemblies of each taxonomic group (Table 2, Supplementary Material 1 - Figure 3).

**Table 2.** Reference sequences after over-representation filtering. Percentages in brackets show the amount of data left compared to the original set (Table 1). Protein data information can be found in the Supplementary Material 1 - Table 5.

|  | Base pairs | # species | # leaf taxids | # assemblies | # sequences |
| --- | --- | --- | --- | --- | --- |
| RefSeq-CG-top-3 | 2.9e10<br>(62%) | 11,464<br>(100%) | 14,071<br>(100%) | 15,171<br>(77%) | 24,290<br>(74%) |
| RefSeq-ALL-top-3 | 2.1e11<br>(36%) | 29,061<br>(100%) | 51,292<br>(100%) | 56,805<br>(38%) | 4,400,402<br>(29%) |

For classification we used reads from the first CAMI Challenge (Sczyrba *et al.*, 2017). Sets of simulated datasets mimicking commonly used environments and settings were obtained, representing multiple closely related strains, plasmids and viral sequences. These samples were divided into 3 categories: low, medium and high complexity with increasing number of organisms and different sequencing profiles providing a well-produced and challenging dataset to analyze. The prechallenge simulated reads were generated based on public data (NCBI, SILVA46) and an exact ground truth assignment is provided for each read down to sequence level. The challenge datasets were more realistic and obtained from newly sequenced genomes of 700 microbial isolates and 600 circular elements and a ground truth is provided at taxonomic levels. Here we used one high complexity sample from both the pre-challenge dataset (simulated) and the realistic challenge dataset (real) categories to perform evaluations and benchmark the tools (Supplementary Material 1 - Table 6).

The classification results were evaluated in terms of sensitivity and precision in two different ways: cumulative- and rank-based. Details on their differences can be found in the Supplementary Material 1 - Section 2.7. In short, the cumulative-based evaluation will compare how well tools perform up to a certain taxonomic level, considering only the taxon of their final classification level. The rank-based evaluation considers the full lineage of each classification. For example: in a cumulative-based evaluation, values of sensitivity and precision at family level will account cumulatively for all sequences classified at subsequent taxonomic levels (genus, species, species+) up to (and including) the family level. In a rank-based evaluation, family level sensitivity and precision values are calculated based on the family assignment from the lineage of the classified sequences. The cumulative-based evaluation provides a better way to compare tools and their ability to correctly classify sequences to their targets. The rank-based approach will better compare how tools perform at a specific taxonomic level. In this work we will use both methodologies to compare the results of the evaluated methods. Additionally, we evaluated all scenarios with AMBER (Meyer *et al.*, 2018), an independent tool for assessment of metagenome binners with a similar approach to the rank-based evaluation. The complete cumulative-based, rank-based and amber results are in the Supplementary Material 2.

The results for the CAMI simulated and real datasets should be interpreted considering the depth of classification. Most tools classify at a certain taxonomic level, either specific rank (e.g. species) or any taxon. Clark provides only species assignments and it was evaluated together with all other tools providing results at any taxonomic level (centrifuge, diamond, ganon, kraken, kraken2). Centrifuge, ganon and krakenuniq are also able to classify sequences at assembly level. Centrifuge outputs at sequence level, thus an extra step of applying an LCA algorithm for non-unique matches was necessary to generate results at assembly and taxonomic levels. Given the availability of the ground truth, only simulated data was evaluated up to assembly level while real data was evaluated at taxonomic levels.

### 3.1 Indexing

The set of reference sequences from RefSeq-OLD/CG/ALL (Table 1) and RefSeq-CG/ALL-top-3 (Table 2) were used as inputs to generate the indices for each evaluated tool. Here evaluation is done by total run-time, memory consumption and final index size (Table 3 and 4).

**Table 3.** Build times, memory consumption and index sizes at taxonomic level. Memory and Index size in GiB. All tools build at taxonomic leaf nodes (taxid) besides clark building at species level. Tools running more than 24 hours to build were not considered. 48 threads were used for all tools. Computer specifications and parameters used are in the Supplementary Material 1 - Section 2.1 and 2.4. Krakenuniq was not evaluated on taxonomic level since it runs exactly the same base algorithm as kraken in this configuration.

| Reference | Method | time | Memory | Index size |
| --- | --- | --- | --- | --- |
| RefSeq-OLD | centrifuge | 02:51:03 | 98 | 4 |
|  | clark | 04:07:56 | 150 | 32 |
|  | diamond | 00:08:07 | 28 | 3 |
|  | ganon | 00:02:08 | 22 | 15 |
|  | kraken | 02:04:16 | 87 | 73 |
|  | kraken2 | 00:17:28 | 13 | 10 |
| RefSeq-CG-top-3 | centrifuge | 06:51:25 | 262 | 12 |
|  | clark | 08:45:31 | 243 | 81 |
|  | diamond | 00:10:33 | 27 | 9 |
|  | ganon | 00:07:01 | 68 | 62 |
|  | kraken | 04:53:31 | 195 | 184 |
|  | kraken2 | 00:45:25 | 29 | 26 |
| RefSeq-ALL-top-3 | diamond | 00:36:23 | 30 | 70 |
|  | ganon | 00:54:48 | 248 | 249 |
|  | kraken2 | 05:04:24 | 124 | 123 |

**Table 4.** Build times, memory consumption and index sizes at assembly level. Memory and Index size in GiB. Tools running more than 24 hours to build were not considered. 48 threads were used for all tools. Computer specifications and parameters used are in the Supplementary Material 1 - Section 2.1 and 2.4

| Reference | Method | time | Memory | Index size |
| --- | --- | --- | --- | --- |
| RefSeq-OLD | centrifuge | 02:51:03 | 98 | 4 |
|  | ganon | 00:02:22 | 30 | 23 |
|  | krakenuniq | 02:06:41 | 87 | 73 |
| RefSeq-CG | centrifuge | 12:32:08 | 428 | 20 |
|  | ganon | 00:10:49 | 100 | 93 |
|  | krakenuniq | 08:54:56 | 321 | 190 |
| RefSeq-ALL | ganon | 02:30:47 | 493 | 501 |

When indexing the RefSeq-CG-top-3 at taxonomic levels (Table 3), the evaluated tools took between 7 minutes and 8 hours, resulting in ganon being the fastest and clark the slowest. We do not consider runs taking more than 24 hours to build indices, given that they clearly do not scale well enough to index high amounts of data and will not be able to keep indices up-to-date in a reasonable amount of time for new data (Supplementary Material 1 - Section 3.2). Ganon shows a significant overall reduction in run-time compared to the other tools besides diamond. However, diamond is the only tool using protein data, accounting approximately for a third of the volume of the genomic data. Ganon builds 6 times faster than kraken2, the second fastest using the same data source. Centrifuge achieves the lowest index size with the cost of having the highest memory consumption. Additionally, ganon is able to generate smaller indices and use less memory at the cost of speed in the classification step, without harming sensitivity (Supplementary Material 1 - Section 3.5). Ganon indices for RefSeq-CG-top-3 can be as small as 21GiB. RefSeq-ALL-top-3 was built in under an hour for diamond and ganon with kraken2 taking more than 5 hours. Diamond generated the smallest filter and achieved the lowest memory consumption. We could not run centrifuge, clark, kraken and krakenuniq for RefSeq-ALL on our infrastructure, given computational limitations or long execution time. A recent publication (Nasko *et al.*, 2018) reported that kraken and consequently krakenuniq both need 11 days to build a database for the bacterial RefSeq version 80, an approximate of the RefSeq-ALL here evaluated, with a more powerful server consisting of 64 cores of E7-8860v4 CPUs and three terabytes of memory. Estimated run times for these tools in the evaluated datasets can be found in the Supplementary Material 1 - Figure 7.

When building indices on assembly level (Table 4), ganon took around 10 minutes to index RefSeq-CG while the second fastest tool, krakenuniq, took almost 9 hours. Given our computational and time limitations, ganon was the only tool able to build indices on assembly level for the RefSeq-ALL dataset, taking 2 hours and 30 minutes.

### 3.2 Updating

Ganon is the only tool among the evaluated ones that allows for incremental updates on previously generated indices. We evaluated this functionality on Bacterial sequences added to RefSeq-CG dating from 19-December-2018 to 21-January-2019, comprising 2.77 Gbp, 1307 sequences, 370 species from which 213 are new to the reference set and 716 new assemblies (Supplementary Material 1 - Table 4). Updating the ganon index based on RefSeq-CG with this dataset finished under 5 minutes, less than half of the time necessary to create the index (Table 4).

### 3.3 Classifying

Figure 2 compares in a cumulative-based fashion the results of one simulated high complexity dataset (CAMI toy set) classified against the indices based on RefSeq-OLD, RefSeq-CG-top-3 and RefSeq-ALL-top-3. In this analysis we can observe how each method performs classifying reads to their ground truth targets up to a certain taxonomic level. The overall improvement in terms of sensitivity and precision is clear when using a more complete and up-to-date set of references (RefSeq-ALL-top-3), since they provide higher coverage for the evaluated ground truth targets (Supplementary Material 1 - Figure 6). The highest F1-Score at any taxonomic level is achieved by ganon and kraken2, both performing similarly using RefSeq-ALL-top-3. Diamond shows an increase in performance at higher taxonomic levels but performs poorly at species level. Clark classifies only at species level and has no improvements in higher taxonomic levels. Metrics for the complete RefSeq-CG and RefSeq-ALL differ slightly from the respective top-3 sets, therefore they were not included in the evaluations (Supplementary Material 1 - Section 3.3.1). This indicates that over-representation filtering does not affect the results but it can be used to speed up analysis.

When looking at the metrics by each rank individually (Table 5, Supplementary Material 1 - Figure 8), the overall precision and sensitivity values are greater, since incorrect classifications at lower levels are not penalized in this type of evaluation. Besides diamond, which underperforms at species level, all tools have overall similar performance values using RefSeq-OLD and RefSeq-CG-top-3. However, tools show improvement on sensitivity on all levels with RefSeq-ALL-top-3. Ganon is 12% more sensitive at species level with this dataset and reaches 99.54% precision at genus level (Supplementary Material 2). For lower ranks (species and species+) results were mainly limited by the availability of reference sequences (Supplementary Material 1 - Figure 6).

**Table 5.**
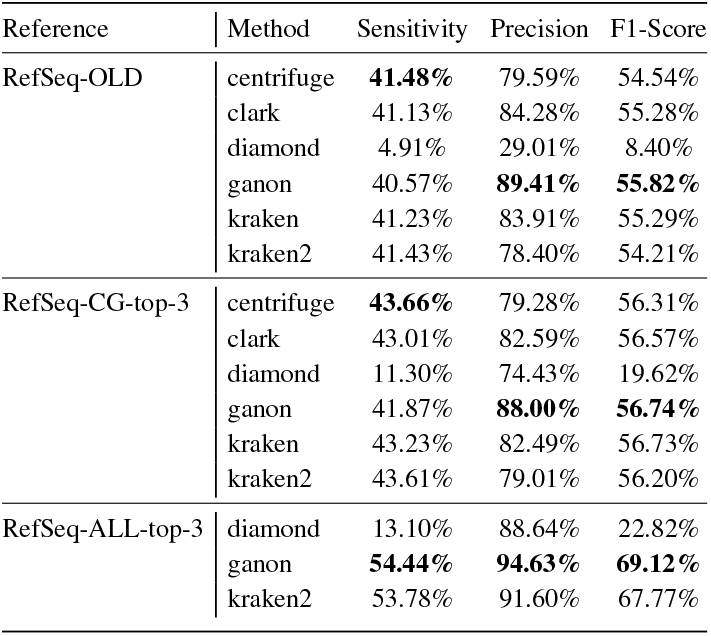
Rank-based precision, sensitivity and F1-Score values for the simulated reads at species level. The use of a larger reference set with RefSeq-ALL-top-3 significantly improves results. Only ganon and diamond indexed the RefSeq-ALL-top-3 in less than 24 hours, thus centrifuge, clark and kraken were excluded. Results for all taxonomic levels are in the Supplementary Material 1 - Figure 8 and Supplementary Material 2

| Reference | Method | Sensitivity | Precision | F1-Score |
| --- | --- | --- | --- | --- |
| RefSeq-OLD | centrifuge | **41.48%** | 79.59% | 54.54% |
|  | clark | 41.13% | 84.28% | 55.28% |
|  | diamond | 4.91% | 29.01% | 8.40% |
|  | ganon | 40.57% | **89.41%** | **55.82%** |
|  | kraken | 41.23% | 83.91% | 55.29% |
|  | kraken2 | 41.43% | 78.40% | 54.21% |
| RefSeq-CG-top-3 | centrifuge | **43.66%** | 79.28% | 56.31% |
|  | clark | 43.01% | 82.59% | 56.57% |
|  | diamond | 11.30% | 74.43% | 19.62% |
|  | ganon | 41.87% | **88.00%** | **56.74%** |
|  | kraken | 43.23% | 82.49% | 56.73% |
|  | kraken2 | 43.61% | 79.01% | 56.20% |
| RefSeq-ALL-top-3 | diamond | 13.10% | 88.64% | 22.82% |
|  | ganon | **54.44%** | **94.63%** | **69.12%** |
|  | kraken2 | 53.78% | 91.60% | 67.77% |

The same analysis was performed on real data (CAMI challenge set). This set is more challenging since most of the species in the sample are novel and, still to this date, mostly not present in the analyzed repositories of reference sequences (Supplementary Material 1 - Figure 6). As stated by the CAMI results (Sczyrba *et al.*, 2017), tools performed poorly in this dataset in terms of sensitivity (Figure 3). Here the impact of a larger and up-to-date set of references (RefSeq-ALL-top-3) is more evident, thus significantly improving the results on both sensitivity and precision. The same trend from the simulated data analysis is present, with ganon and kraken2 achieving best results up to species level. Diamond improves classifications at higher levels but has poor resolution at lower ranks, showing a very conservative approach. Diamond classifies more reads at lower taxonomic ranks (family and above) than the other methods. However, on a rank-based evaluation diamond is overall less sensitive than ganon and kraken2.

**Fig. 3.**
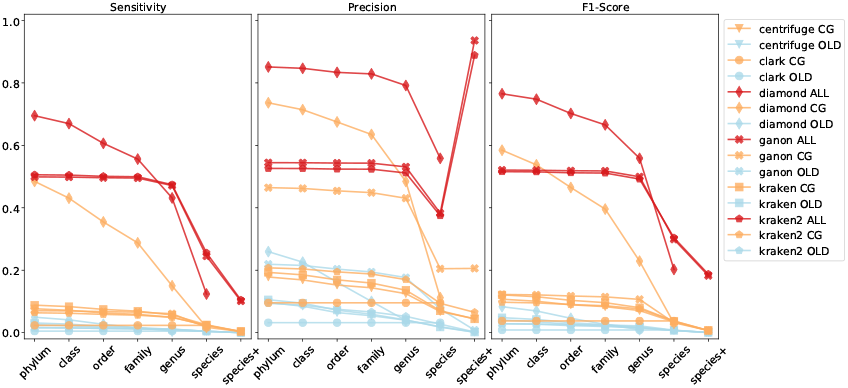
Cumulative-based precision, sensitivity and F1-Score values at all ranks for the real reads against all evaluated reference sets (blue = RefSeq-OLD, orange = RefSeq-CG-top-3, red = RefSeq-ALL-top-3)

In the rank-based analysis (Table 6, Supplementary Material 1 - Figure 9) ganon and kraken2 have 10% higher F1-Score compared to diamond with the RefSeq-ALL-top-3 at species level. Sensitivity has a peak of 10% and 25% at species+ and species levels, respectively, which are not far from the maximum possible using this reference set (12% and 32% respectively). The high peak in precision at species+ level can be explained by the reduced number of reads in the ground truth assigned to taxonomic entries of those ranks (e.g. subspecies), being 17% of the real reads and 63% of the simulated reads. In addition, fewer reads are classified uniquely among similar genomes at this level, resulting in a higher level LCA classification. Comparing results between RefSeq-CG-top-3 and RefSeq-ALL-top-3, genus level sensitivity went from 13% to 83% with a significant improvement in precision, reinforcing the need for bigger and more diverse reference sets to analyze metagenomics data. Similar results can be seen in amber evaluation (Figure 4).

**Table 6.** Rank-based precision, sensitivity and F1-Score values for the real reads at species level. The use of a larger reference set with RefSeq-ALL-top-3 significantly improves results. Only ganon and diamond indexed the RefSeq-ALL-top-3 in less than 24 hours, thus centrifuge, clark and kraken were excluded. Results for all taxonomic levels are in the Supplementary Material 1 - Figure 9 and Supplementary Material 2

| Reference | Method | Sensitivity | Precision | F1-Score |
| --- | --- | --- | --- | --- |
| RefSeq-OLD | centrifuge | 0.51% | 2.24% | 0.84% |
|  | clark | 0.49% | 3.21% | 0.86% |
|  | diamond | 0.00% | 0.00% | 0.00% |
|  | ganon | 0.43% | **7.75%** | 0.82% |
|  | kraken | 0.50% | 3.13% | 0.86% |
|  | kraken2 | **0.55%** | 2.19% | **0.87%** |
| RefSeq-CG-top-3 | centrifuge | 2.41% | 7.03% | 3.59% |
|  | clark | 2.34% | 9.57% | 3.76% |
|  | diamond | 1.74% | 11.23% | 3.02% |
|  | ganon | 1.77% | **0.89%** | 3.26% |
|  | kraken | 2.39% | 9.61% | **3.83%** |
|  | kraken2 | **2.59%** | 7.22% | 3.82% |
| RefSeq-ALL-top-3 | diamond | 12.38% | **55.84%** | 20.27% |
|  | ganon | 24.78% | 38.67% | 30.20% |
|  | kraken2 | **25.83%** | 37.84% | **30.70%** |

**Fig. 4.**
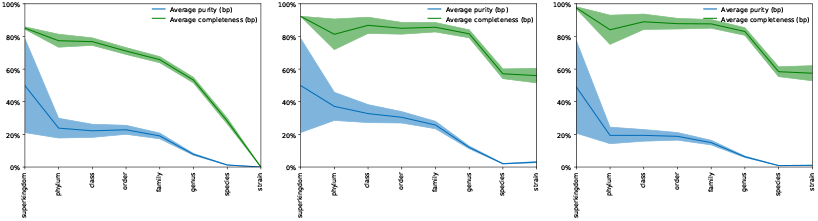
AMBER average completeness/sensitivity (green) andpurity/precision (blue) values for real reads. Results for diamond (left), ganon (middle) and kraken2 (right) using RefSeq-ALL-top-3 set of references. Strain level in AMBER plots are equivalent to species+ in our evaluations.

Table 7 compares the assembly level classification between centrifuge, ganon and krakenuniq. There is an overall decrease in precision and sensitivity from RefSeq-OLD to RefSeq-CG. Precision is greater using RefSeq-ALL but sensitivity is still greater with RefSeq-OLD. However, RefSeq-CG has more than 6 times the number of assemblies of RefSeq-OLD, while RefSeq-ALL has almost 50 times more assemblies (Table 1). As reported before (Nasko *et al.*, 2018), higher diversity in the references does not always translate to an improved accuracy in the classification. This was also noticed when using the complete NCBI-nt database to analyze the same dataset (Supplementary Material 1 - Figure 14).

**Table 7.** Rank-based precision, sensitivity and F1-Score values for the simulated reads at assembly level. Only ganon indexed the RefSeq-ALL in less than 24 hours, thus centrifuge, clark and kraken were excluded. Results for all taxonomic levels are in the Supplementary Material 2.

| Reference | Method | Sensitivity | Precision | F1-Score |
| --- | --- | --- | --- | --- |
| RefSeq-OLD | centrifuge | **22.78%** | 64.54% | 33.68% |
|  | ganon | 22.32% | **77.95%** | **34.70%** |
|  | krakenuniq | 22.68% | 69.66% | 34.22% |
| RefSeq-CG | centrifuge | **11.82%** | 30.77% | 17.08% |
|  | ganon | 11.52% | **37.25%** | **17.60%** |
|  | krakenuniq | 11.67% | 32.45% | 17.17% |
| RefSeq-ALL | ganon | 21.56% | 87.89% | 34.62% |

In the specific case of methods evaluated here, small differences between very similar assemblies are difficult to be identified due to the resolution of each method. This means that they, in general, can correctly classify sequences to target assemblies given a certain similarity threshold. However, they are unable to select the correct assembly, thus providing the lowest common ancestor at a lower resolution. This can be seen in Supplementary Material 1 - Figure 13, where the overall sensitivity and precision of all tools executing in assembly mode did not affect the taxonomic metrics and are comparable to the same tools running in taxonomic mode. Even though the assembly step does not provide accurate enough results, centrifuge and ganon are the only tools that can provide a list of all matches/candidates that can be further analyzed with high resolution methods (Fischer *et al.*, 2017).

In most scenarios evaluated, ganon consistently provides greater precision classifying reads to their ground truth targets within the same reference set, while keeping sensitivity values high, with little variation to the other methods. High precision translates to fewer reads with a wrong classification. Sensitivity is strongly improved in more diverse reference sets, especially with RefSeq-ALL-top-3. Looking at rank-by-rank performance, ganon and kraken2 improved F1-Score in every taxonomic rank (Supplementary Material 1 - Figures 8 and 9), with F1-Score up to 46% higher than diamond with the same reference at species level (Table 5).

Table 8 compares the performance of the analyzed tools in terms of how many base pairs they can classify per minute (Mbp/m), wall/elapsed time and memory usage. Kraken2 is the tool with the fastest runtime on classification step and diamond with the slowest. Although comparisons with diamond were made, it is important to notice that the tool works in a very different way using protein data and performing alignments, thus explaining the huge difference in execution times and results. Ganon can be configured to run in *off set* mode, thus skipping a certain number of k-mers and speeding up classification. *off set* = 1 means that all k-mers are being evaluated while *off set* = 2 means that every 2nd k-mer is being skipped. The trade-off between offset and precision/sensitivity for ganon results can be seen in Supplementary Material 1 - Figure 12. Speed variation between simulated and real reads is partly explained due to their classification rate: on average 70% of the simulated reads are classified while only 20% of the real reads are classified. Memory consumption is mainly based on the index size of each tool (Table 3), with little variation besides that.

**Table 8.** Classification performance. Memory in GiB. Full set of simulated and real reads classified with 48 threads. Centrifuge, clark and diamond performance in Mbp/m calculated from wall time. Values are the average of 4 out 5 consecutive runs (excluding the slowest run), with standard deviation for the run time in parentheses. Computer specifications and parameters used are in the Supplementary Material 1 - Section 2.1 and 2.4

| Reference | Method | Simulated |  |  | Real |  |  |
| --- | --- | --- | --- | --- | --- | --- | --- |
|  |  | Mbp/m | Wall time | Memory | Mbp/m | Wall time | Memory |
| RefSeq-CG-top-3 | centrifuge | 298 | 00:24:59 (± 51s) | 13 | 802 | 00:09:19 (± 4s) | 13 |
| | clark | 1104 | 00:06:44 (± 5s) | 101 | 1208 | 00:06:11 (± 4s) | 100 |
| | diamond | 36 | 03:27:00 (± 259s) | 14 | 33 | 03:40:55 (± 170s) | 15 |
| | ganon | 380 | 00:20:31 (± 8s) | 61 | 538 | 00:14:50 (± 1s) | 61 |
| | kraken | 2113 | 00:03:46 (± 1s) | 177 | 2734 | 00:02:57 (± 3s) | 177 |
| | kraken2 | 3085 | 00:02:47 (± 1s) | 27 | 3833 | 00:02:19 (± 1s) | 27 |
| RefSeq-ALL-top-3 | diamond | 6 | 18:23:09 (± 729) | 21 | 5 | 21:23:00 (± 181s) | 22 |
| | ganon | 107 | 01:13:44 (± 7s) | 243 | 153 | 00:53:13 (± 12s) | 243 |
| | kraken2 | 2991 | 00:04:11 (± 4s) | 123 | 3659 | 00:03:52 (± 1s) | 122 |

## 4 Discussion

We presented ganon, a novel method to index big sets of genomic sequences and classify short reads against them in a taxonomic oriented scheme. Ganon’s strengths are an ultra-fast indexing method for large sets of reference sequences that incorporates a novel application of Interleaved Bloom Filters and aprecise classification with k-mer counting and filtering. Unlike DREAM-Yara, an alignment-based read mapper that uses the IBF as a pre-filter for the distributed Yara mapper, ganon uses the IBF as the main index structure to provide an alignment-free assignment of sequences. This is only possible by creating taxonomic constrained clusters with TaxSBP in any desired taxonomic level. Ganon additionally applies an LCA algorithm as a final step to have one classification per sequence. In addition it also provides updatability of indices, multi-hierarchy support for classification, assembly level support and taxonomic reports.

By indexing large sets of reference sequences and turn them into searchable indices, ganon allows scientists to make most of their data. Short turnaround times for index building and updating are crucial for many bioinformatics applications (e.g. outbreak investigation). In our evaluations, building the complete RefSeq and classifying 49 million reads against it performed under 2 hours with ganon, from raw reference sequences and reads to taxonomic reports, while kraken2 required more than 5 hours and diamond more than 22 hours to index and classify the same set. Other methods required at least 24 hours to build the indices. Without a dedicated infrastructure for constant reconstruction of indices and databases, tools evaluated in this work are unable to keep up with the fast growing rate of reference sequence repositories. That results in either long time to start analysis or use of outdated reference sets. Ganon facilitates database maintenance, allowing short increments on a daily basis being a realistic option to keep-up with the fast pace of data generation. In addition, ganon indices are flexible and can be built for different taxonomic levels (e.g. genus), requiring less space and memory, consequently improving classification speed. A trade-off between filter size, clustering and false positive rate is also possible, simply by sacrificing precision over performance or disk usage over classification speed (Supplementary Material 1 - Section 3.5).

Classification results presented here are on par with state-of-the-art methods with regards to sensitivity, while improving precision rates in almost every scenario of our cumulative-based evaluations. Results are consistent across all three evaluation methods (cumulative- and rank-based and amber) indicating the robustness of findings. We attribute this improvement to an application of the k-mer counting lemma together with a progressive filtering step, which can better separate false from true positives. The unique filtering step also allows for better selection of false positives when taxonomic groups are underrepresented in the reference set. In addition, instead of only reporting reads at a fixed LCA level, ganon provides every output for a read at a taxonomic or assembly level. This is crucial for strain level analysis, where candidate organisms are more insightful for further investigations than a conservative identification.

Even with ganon achieving improved results in classification, in general terms, the methods tested here perform similarly when based on the same underlying set of reference sequences. The difference in sensitivity when using a high quality set (RefSeq-ALL) compared to only complete genomes (RefSeq-CG) or an outdated set (RefSeq-OLD) is very significant and tends to get bigger with more sequences added to this repository. Thus the choice of the database is crucial and should not be overlooked when analyzing metagenomics data. Even though centrifuge, clark, kraken and krakenuniq could potentially perform well with more reference sequences, their indexing times are highly prohibitive.

When using highly diverse reference sets or when aiming at high resolution classification (e.g. assembly level), the evaluated methods shown decreased performance. However, in a scenario of data exploration of an unknown environmental sample, the ability to classify reads against huge sets of very diverse reference sequences (e.g. NCBI-nt) can be helpful. Therefore, in those scenarios we recommend to perform analysis hierarchically, first classifying reads against high quality references and only using high diverse reference sets for unclassified sequences, adjusting error rates accordingly. This approach can be easily done with ganon’s implementation of multi-filter and multi-hierarchy classification. This functionality tied to fast indexing of reference sets make ganon a powerful tool for exploratory data analysis, enabling multiple combinations of indices and error rates in an iterative manner. An example of this functionality can be found in Supplementary Material 1 - Section 3.4, where we analyzed real data from TARA oceans (Tully *et al.*, 2018), building several indices and classifying reads against them in an exploratory-fashion.

Ganon’s fast indexing performance is mainly due to the fact that k-mers are not being counted. Instead, all of them are inserted into a space-efficient data structure (IBF) that also provides quick look-up times. However, data generation is constantly increasing and in the long term this approach will reach a limit. For that reason, a k-mer aware clustering combined with a minimizer implementation could improve performance in the data structure as well as memory consumption. These features are planned for future releases. Even though we based our analysis on large and realistic datasets, time efficiency purely based on data can be misleading. Thus, the scalability of the methods can only be deduced. As a future work we propose a comparison of time and space complexities of each methodology and how they would perform in the long term, considering a continuous and exponential data growth.

Ganon manages to index large sets of reference sequences while keeping them updated in very short time. In addition, classification results for ganon are as good as or better than the evaluated tools and it runs in competitive time. To the best of our knowledge, ganon is the only tool with update capabilities, which is performed in a fraction of the complete build time. This poses as an advantage to maintain up to date with the public repositories of genomic data and their frequent updates. To conclude, we believe that ganon can be a useful tool for metagenomics analysis in a time where reference sequence repositories are growing fast.

## Acknowledgements

We would like to thank Tobias Loka for helpful discussions, Diogo Andrei Benvenutti and Fabio Malcher Miranda for git project improvements and C++ contributions, Robert Rentzsch and Elizabeth Yuu for revising the manuscript.

