## Supplementary Material 1 for "ganon: precise metagenomics classification against large and up-to-date sets of reference sequences"

### 1 Introduction

Figure 1 shows the amount of reference sequences available over the last 11 years in the GenBank [1] and RefSeq [2] repositories from NCBI. The growth is exponential. RefSeq sequences from Archaeal and Bacterial genomes are highlighted for being a commonly used reference set for classification in metagenomics.

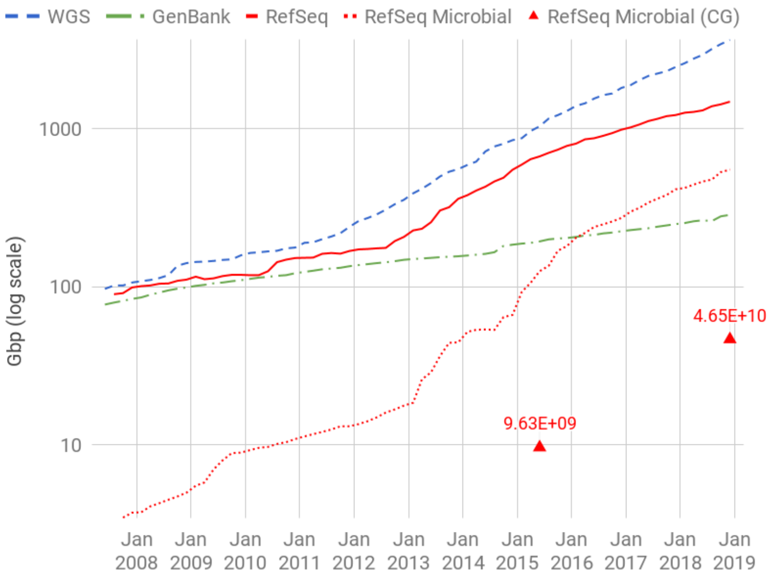

Figure 1: **Number of available sequences in NCBI repositories from June 2007 to December 2018 on a logarithmic scale.** Microbial stands for Archaeal and Bacterial organisms and CG stands for Complete Genomes. RefSeq Microbial has an uninterrupted and linear growth on a logarithmic scale. Data collected from: https://ftp.ncbi.nlm.nih.gov/refseq/release/release-statistics/ and https://www.ncbi.nlm.nih.gov/genbank/statistics/

### 2 Methods

#### 2.1 Computer Specifications

Results presented in the manuscript were executed in a virtual machine provided by denbi cloud (https://www.denbi.de/cloud) with the following specifications:

- 56 VCPUs - Intel Xeon Processor (Skylake, IBRS), 2593 MHz
- 1968 GiB RAM
- Linux version 4.15.0-45-generic
- Ubuntu 18.04.1 LTS
- gcc version 7.3.0 (Ubuntu 7.3.0-16ubuntu3)

#### 2.2 Time Complexity

In the construction of the Interleaved Bloom Filter, we compute a constant number $h$ of hash functions (3 by default) in $O(1)$ for each k-mer in the text of length $n$. Therefore, the construction step runs in $O(n)$.

To search a query, we take a $h$ sub-bit vectors of length $n$ and $AND$ it. This takes $O(b/w)$ time where $b$ is the number of bins and $w$ the word length (i.e. 64) and assuming a constant $h$. Then we increment counters for each 1 in the "anded" bitvector. If the k-mer occurs in $x$ bins, this takes time $O(x)$ (using lzcount). Hence the runtime is $O(b/w + x)$. This can be assumed as constant is for many cases (i.e. $b = 256$, $w = 64$, $x = 2$).

#### 2.3 Tools version

All results, evaluation and plots presented in the main manuscript were compiled in a Snakemake [3] benchmark pipeline for easy validation and reproducibility and can be found at: https://github.com/pirovc/ganon_benchmark.

The version of the pipeline used to generate the results presented is 1.1.0.

All evaluated tools, besides AMBER which should be provided externally, are available in the Bioconda repository [4] and are automatically installed when running the pipeline with the useconda parameter in the following versions:

- AMBER=2.0.21-beta
  - numpy=1.16.4
  - biopython=1.73.0
  - matplotlib=3.1.1
  - bokeh=0.13.0
  - pandas=0.24.2
  - seaborn=0.9.0
- centrifuge=1.0.3
- clark=1.2.5
- diamond=0.9.24
- ganon=0.2.1
- krakenuniq=0.5.5
- kraken=1.0
  - jellyfish=1.1.12
- kraken2=2.0.8-beta

#### 2.4 Tools parameters

The exact commands executed for each tool can be found on the ganon_benchmark pipeline versions 1.0.0. The most important parameters used for each tool different than the default are listed below:

##### 2.4.1 Ganon parameters

Ganon deafult parameters were used to build references, as follows:

- k-mer size: 19
- Number of hash functions: 3
- Max. false positive: 0.05

##### 2.4.2 Build

- RefSeq-CG
  - diamond: taxonmap prot.accession2taxid.gz taxonnodes nodes.dmp
  - ganon (taxid): -r taxid -m 96000
  - ganon (assembly): -r assembly -m 96000
  - krakenuniq: taxids-for-genomes
- RefSeq-CG-top-3
  - diamond: taxonmap prot.accession2taxid.gz taxonnodes nodes.dmp

  - ganon: -r taxid -m 64000

- RefSeq-ALL
  - diamond: taxonmap prot.accession2taxid.gz taxonnodes nodes.dmp
  - ganon (taxid): -r taxid -m 480000
  - ganon (assembly): -r assembly -m 512000

- RefSeq-ALL-top-3
  - diamond: taxonmap prot.accession2taxid.gz taxonnodes nodes.dmp
  - ganon: -r taxid -m 256000

###### 2.4.3 Classify

- Simulated
  - ganon: max-error 3 max-error-unique 2 offset 2

- Real
  - ganon: max-error 5 max-error-unique 4 offset 3

##### 2.5 Data

###### 2.5.1 References

Detailed information on the set of reference sets used in the evaluations is provided below: RefSeq-OLD (Table 1), RefSeq-CG (Table 2), RefSeq-ALL (Table 3) and update data (Table 4). The set of reference sequences used for diamond are detailed in the Table 5. All reference data, besides RefSeq-OLD, was downloaded using https://github.com/pirovc/genome_updater.

| | bacteria |
|---|---|
| base pairs | 9,632,441,987 |
| # sequences | 5,242 |
| # species | 1,526 |
| # leaf nodes (taxid) | 2,749 |
| # assembly | 3,042 |

Table 1: **RefSeq-OLD** RefSeq entries of complete genomes of bacteria from 02-June-2015

| | total | archaea | bacteria | fungi | viral |
|---|---|---|---|---|---|
| base pairs | 46,986,899,184 | 699,198,411 | 45,837,996,120 | 191,815,549 | 257,889,104 |
| # sequences | 33,029 | 415 | 22,447 | 85 | 10,082 |
| # species | 11,464 | 212 | 3,638 | 9 | 7,605 |
| # leaf nodes (taxid) | 14,071 | 251 | 5,979 | 9 | 7,832 |
| # assembly | 19,623 | 275 | 11,485 | 9 | 7,854 |

Table 2: **RefSeq-CG** RefSeq entries of complete genomes of archaea, bacteria, fungi and viral groups from 19-December-2018

###### 2.5.2 Reads

The first high complexity sample from the pre-challenge (simulated) and challenge (real) reads (Table 6) were used in this work to perform all evaluations.

| | total | archaea | bacteria | fungi | viral |
|---|---|---|---|---|---|
| base pairs | 587,607,072,429 | 2,360,671,191 | 576,814,972,143 | 8,166,179,365 | 265,249,730 |
| # sequences | 15,201,658 | 44,930 | 15,099,935 | 46,564 | 10,229 |
| # species | 29,061 | 513 | 20,565 | 278 | 7,705 |
| # leaf nodes (taxid) | 51,292 | 608 | 42,471 | 281 | 7,932 |
| # assembly | 147,713 | 795 | 138,683 | 281 | 7,954 |

Table 3: **RefSeq-ALL** Complete RefSeq entries of archaea, bacteria, fungi and viral groups from 19-December-2018

| | bacteria |
|---|---|
| base pairs | 2,771,508,930 |
| # sequences | 1,307 |
| # species / new | 370 / 213 |
| # leaf nodes (taxid) / new | 370 / 213 |
| # assembly / new | 716 / 716 |

Table 4: **RefSeq Update** Updated RefSeq entries of complete genomes of bacteria between 19-December-2018 and 21-January-2019

| | Base pairs (AA) | # sequences |
|---|---|---|
| RefSeq-OLD | 2,743,116,230 | 8,723,327 |
| RefSeq-CG | 13,225,608,527 | 42,240,689 |
| RefSeq-ALL | 165,680,470,950 | 539,052,163 |
| RefSeq-CG-top-3 | 8,200,223,442 (62%) | 25,833,206 (61%) |
| RefSeq-ALL-top-3 | 58,463,424,830 (35%) | 185,134,394 (34%) |

Table 5: **Protein data of reference sequences used for evaluations with DIAMOND.** Percentages in brackets show amount of data left compared to the original set.

| | sample | avg. read length | # reads |
|---|---|---|---|
| CAMI toy (simulated) | H_S001__insert_180 | 100bp | 74,557,111 |
| CAMI challenge (real) | RH_S001__insert_270 | 150bp | 49,905,935 |

Table 6: **Read sets used from the CAMI challenge**

#### 2.6 Distribution of reference sequences

The Figures 2 and 3 show the distribution of sequences in the reference sets before and after overrepresentation filtering.

#### 2.7 Classification evaluation

The classification evaluation was performed in a binary fashion. Every read has an assembly or taxonomic assignment defined by the ground truth. We proposed here two ways of defining true and false positives values based on the taxonomic tree and how to account for them in terms of sensitivity and precision.

Most commonly, taxonomic sequence classification is evaluated in a rank-based approach (Figure 4). Every sequence is assigned to a node in the taxonomic tree and the lineage of such assignment is compared to the lineage of the ground truth assignment for the sequence. For every taxonomic level the nodes between both lineages are compared and marked as true positives when matching and false positives when not matching. Consequently, sensitivity and precision of assignments can be calculated directly for each taxonomic level, accounting for the true and false positive values previously calculated.

We propose a slightly different approach to evaluate taxonomic classification, called cumulative-based approach (Figure 5). Here, every sequence classified to a node in the taxonomic tree is

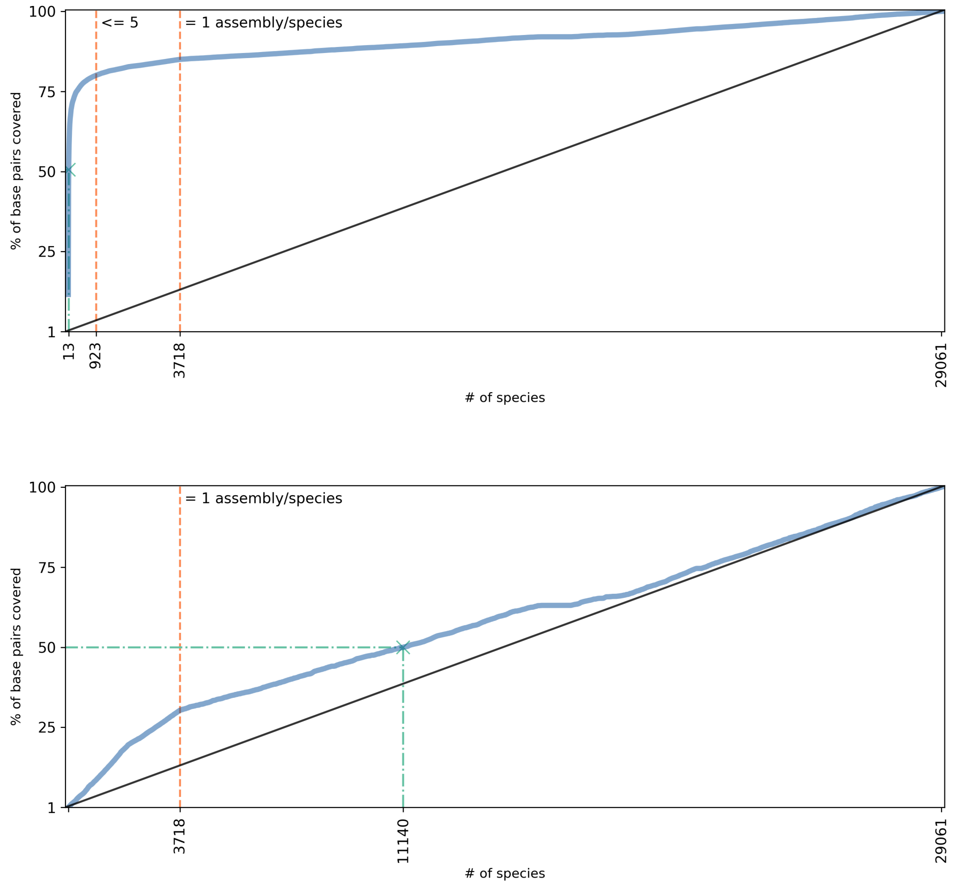

Figure 2: Cumulative sum of base pairs on RefSeq-ALL by species before (top) and after (bottom) the overrepresentation reduction keeping the top 3 biggest assemblies for each species

compared to the leaf node of the ground truth for that sequence. If the assigned node is the leaf ground truth node, it is a true positive. In case a sequence is classified at a higher taxonomic rank than the ground truth leaf node (as long as they have the LCA equal to the ground truth leaf node), it will be considered a false positive - meaning that the sequence is over-classified in the taxonomic tree. When a sequence is classified in the lineage of the ground truth, in other words, when the LCA between the classification node and the ground truth leaf node is the classification node itself, it is a true positive. Finally, when the LCA between the classification node and the ground truth leaf node is any other node of the tree, it will be a false positive. In this approach, every assignment is classified as true or false positive only once, at the taxonomic level of its classification. Sensitivity and precision can be calculated cumulatively from bottom-up of the taxonomic tree, accounting for true and false positives of that level and all higher levels.

Both comparisons are valid approaches to compare the performance of tools in terms of sensitivity and precision for taxonomic sequence classification. However, they differ in some points and should be interpreted differently. The cumulative-based approach will be more sensitive to false assignments, since it will only account for true positives when they are on the lineage of the ground truth. Looking at a certain taxonomic level, the cumulative-based approach will tell how sensitive or precise assignments up to that level are. In a rank-based approach, sensitivity and precision are relative only to the specific level being evaluated. Metrics on a rank-based approach can be misleading on higher taxonomic levels since they do not penalize false assignments on lower taxonomic levels. A cumulative-based evaluation is more effective in comparing how correctly tools are classifying sequences to their truth targets up to a certain taxonomic level while the rank-based approach will evaluate better how well tools perform in a specific taxonomic level independently.

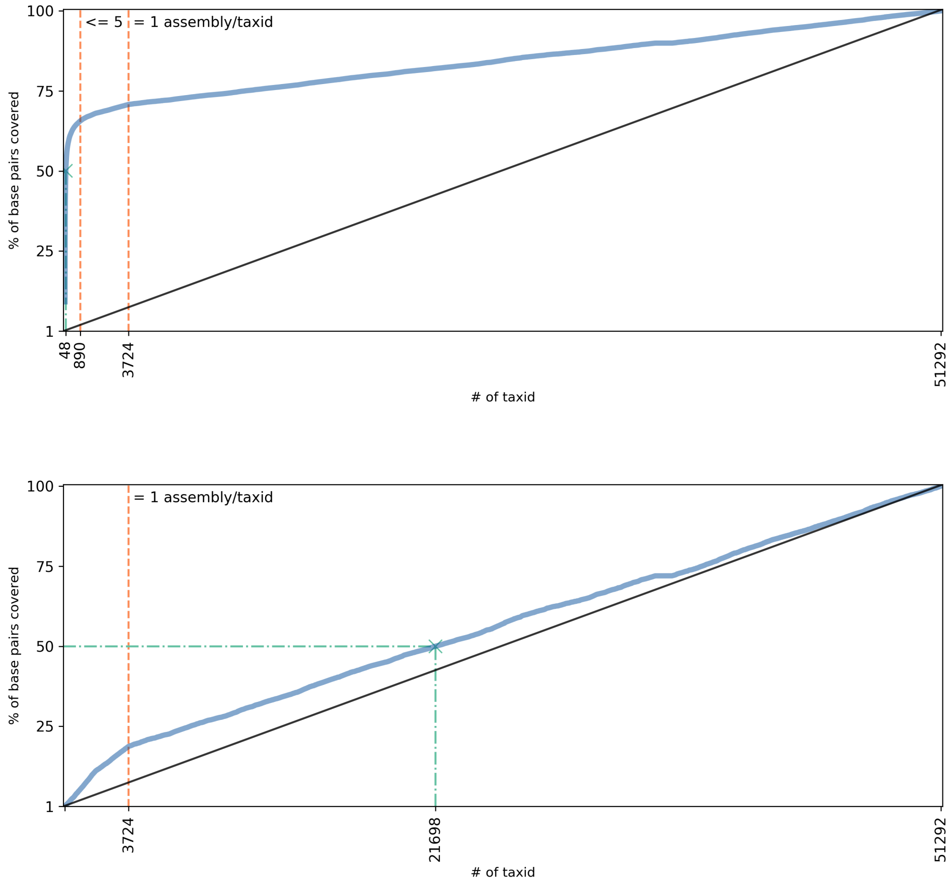

Figure 3: Cumulative sum of base pairs on RefSeq-ALL by taxonomic leaf nodes before (top) and after (bottom) the overrepresentation reduction keeping the top 3 biggest assemblies for each taxid

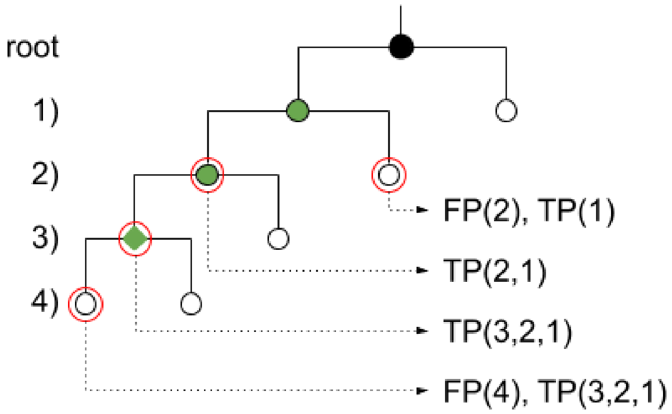

Figure 4: Rank-base evaluation. Taxonomic tree with root node (black) and 4 taxonomic levels (1,2,3,4). Numbers in parentheses after TP and FP represent taxonomic levels. Green nodes represent the ground truth lineage for the green diamond leaf node. Red circles represent all possible classifications in the tree with dotted lines showing true positive and false positive outcomes for each classification.

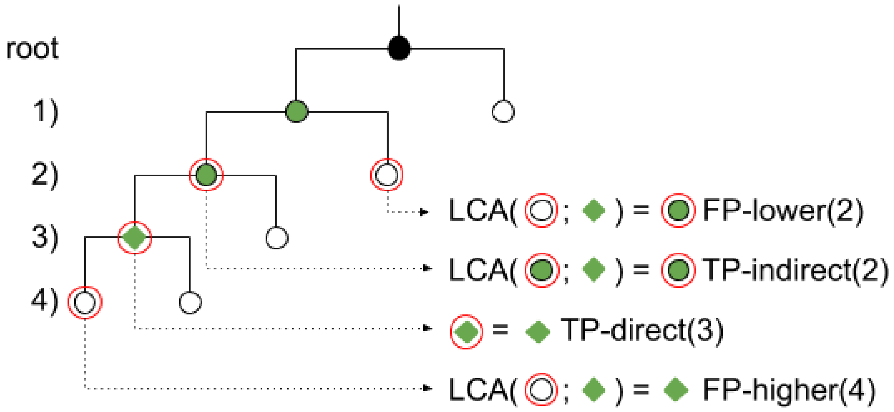

Figure 5: Cumulative-based evaluation. Taxonomic tree with root node (black) and 4 taxonomic levels (1,2,3,4). Numbers in parentheses after TP and FP represent taxonomic levels. Green nodes represent the ground truth lineage for the green diamond leaf node. Red circles represent all possible classifications in the tree with dotted lines showing true positive and false positive outcomes for each classification.

### 3 Results

#### 3.1 Database coverage

Figure 6 shows how many reads could be classified at each taxonomic level based on their ground truth assignments for the reference sets RefSeq-OLD, RefSeq-CG and RefSeq-ALL. Additionally, we compared to the complete NCBI-nt, which was not evaluated in this work, from 01-March-2019.

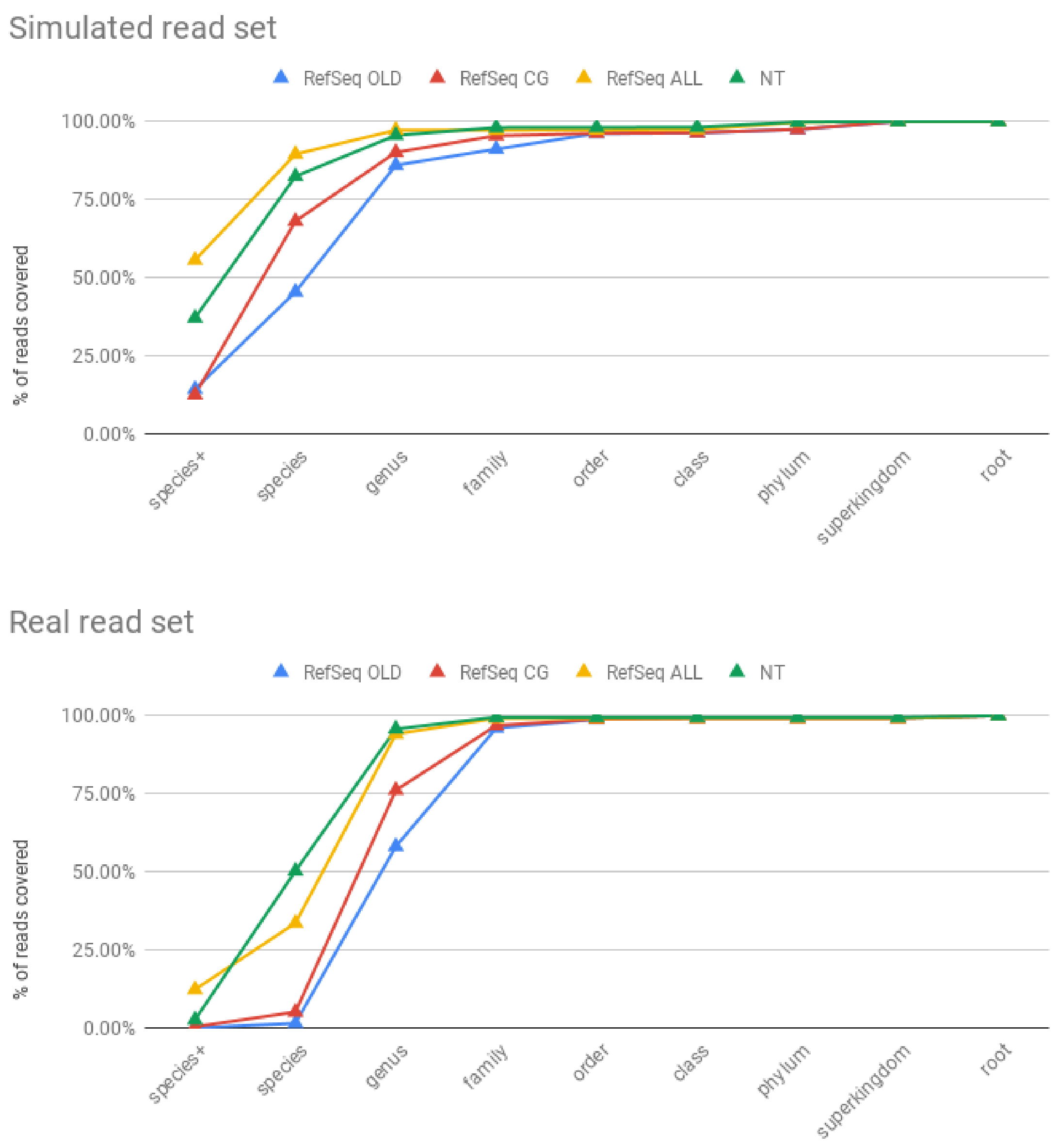

Figure 6: Percentage of simulated (top) and real (bottom) reads with ground truth targets present in the reference sets used + NT

#### 3.2 Indexing

Figure 7 shows actual run times for the indexing step presented in the manuscript with additional estimated linear trend for tools which we did not manage to run in our computational environment (Section 2.1). Once a tool takes more than 24 hours to index, we exclude it from the evaluations.

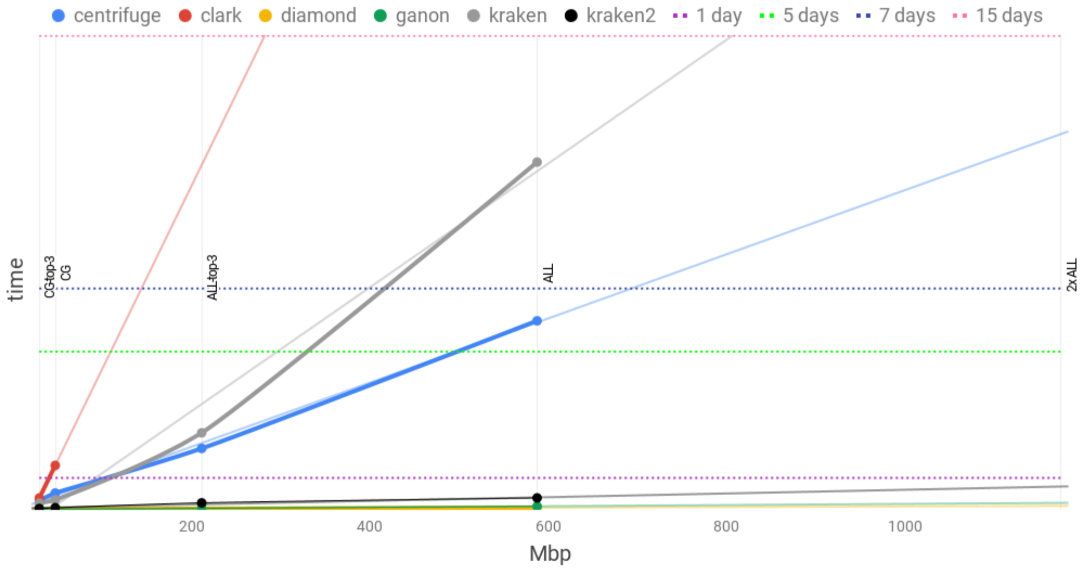

Figure 7: Indexing times (circles). x-axis represents genomic data size in Mbp. Linear trend projection for each tool (same color) in thin lines. 2x RefSeq-ALL was added to simulate a bigger dataset, doubling the size of the biggest reference set evaluated. diamond uses protein data which account, approximately, for a third of the volume of the genomic data.

##### 3.2.1 Ganon indices

The Table 7 shows detailed information on the ganon indices used in the evaluations described in this manuscript. Bin length is automatically calculated to reach a defined filter size (approximately), given a k-mer size, number of hash functions and max false positive rate (default 19, 3 and 0.05 respectively). Fragment length is equal bin length less overlap length (default 300).

| Reference set | Rank | hash | k | bins | bin len. | frag/overlap len. | unique/split | size |
|---|---|---|---|---|---|---|---|---|
| RefSeq-OLD | taxid | 3 | 19 | 2909 | 7004854 | 7004554/300 | 2599/150 | 15 |
| RefSeq-OLD | assembly | 3 | 19 | 3059 | 9994907 | 9994607/300 | 3025/17 | 23 |
| RefSeq-CG-top-3 | taxid | 3 | 19 | 17153 | 4789463 | 4789163/300 | 11730/2341 | 62 |
| RefSeq-CG | assembly | 3 | 19 | 21165 | 5829344 | 5829044/300 | 18117/1506 | 93 |
| RefSeq-ALL-top-3 | taxid | 3 | 19 | 84823 | 3859697 | 3859397/300 | 25979/25311 | 249 |
| RefSeq-ALL | assembly | 3 | 19 | 757931 | 870044 | 869744/300 | 8402/139309 | 501 |

Table 7: **Detailed ganon indices information**. Number of unique/split targets shows how many targets were split among bins (split) and how many are in one bin only (unique). bin len, frag./overlap len in bp. size in GiB.

#### 3.3 Classifying

Rank-based metrics can be found in Figure 8 for the simulated reads and Figure 9 for the real reads.

##### 3.3.1 RefSeq complete and RefSeq top 3

The reference sequences obtained were pre-processed to reduce overrepresentation into "top 3" sets, where only the top 3 biggest assemblies were kept for each taxonomic entry of the set. With such approach, we managed to reduce the data volume to be indexed, keeping the same diversity in the sample. Such reduction did not affect the classification results, as depicted in Figures 10 and 11.

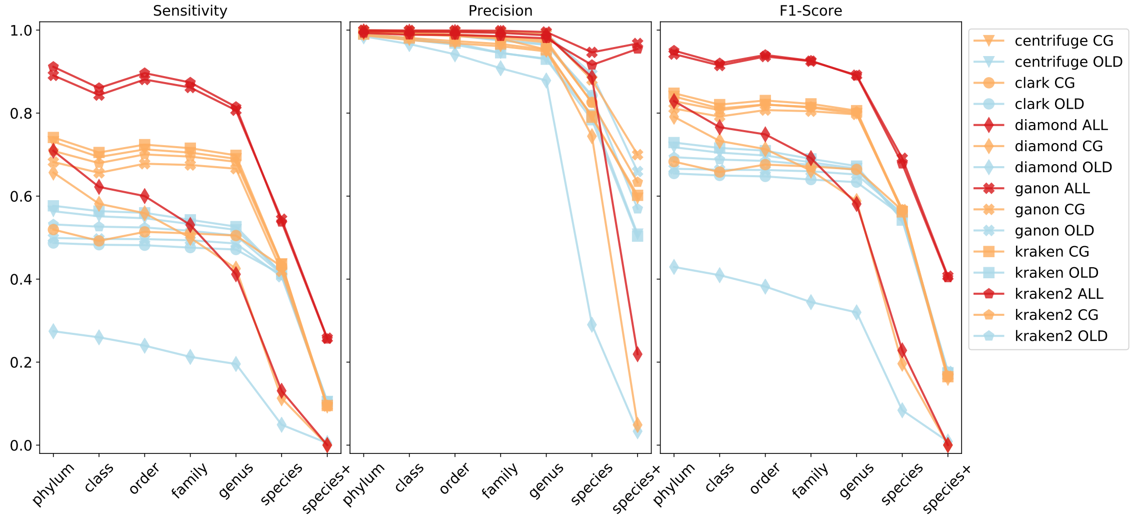

Figure 8: **Rank-based precision, sensitivity and F1-Score for the simulated reads.** Colors represent different reference sets: blue = RefSeq-OLD, orange = RefSeq-CG-top-3, red = RefSeq-ALL-top-3

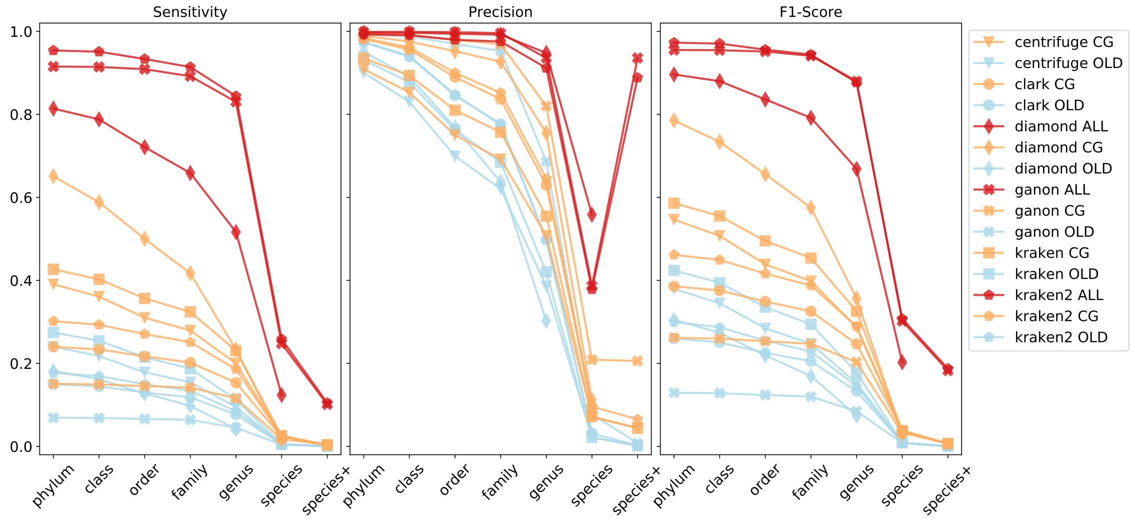

Figure 9: **Rank-based precision, sensitivity and F1-Score for the real reads.** Colors represent different reference sets: blue = RefSeq-OLD, orange = RefSeq-CG-top-3, red = RefSeq-ALL-top-3

##### 3.3.2 Offset

Ganon classification can be executed with an offset $o$, meaning the every $o$-th k-mer will be skipped, speeding up classification mainly at cost of precision. Difference in rank-based metrics from $o = 1..5$ are depicted in Figure 12 against the references from RefSeq-CG-top-3.

##### 3.3.3 Assembly

Figure 13 shows rank-based results for tools classifying at assembly level. The cumulative-based evaluation cannot be applied when classifying at assembly level since it does not follow the same order as the taxonomic classification (e.g. an assembly can be connected to any node of the taxonomic tree).

##### 3.3.4 NCBI-nt

Figure 14 shows rank-based results with the addition of the NCBI-nt database.

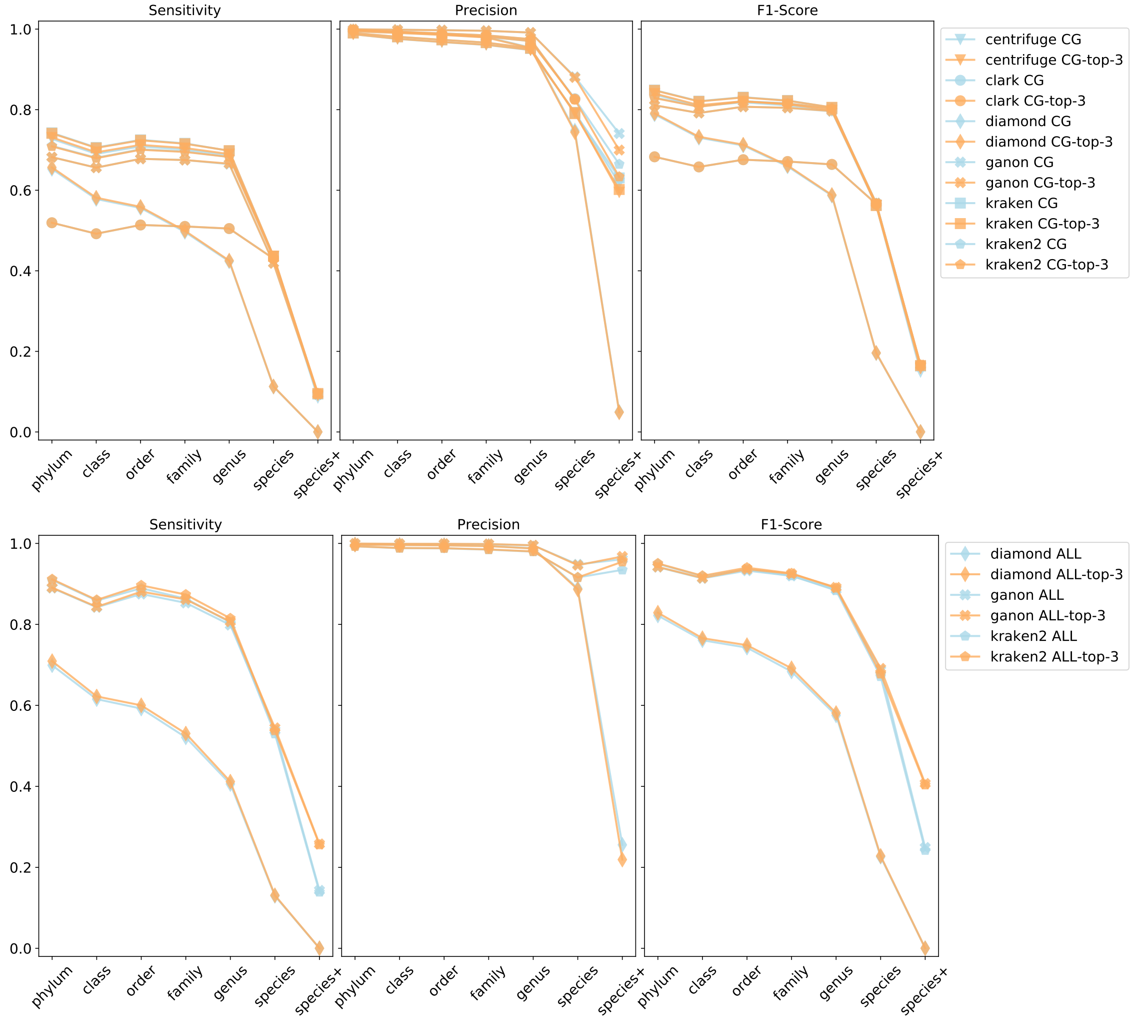

Figure 10: Rank-based precision, sensitivity and F1-Score for the simulated reads between complete RefSeq-CG and RefSeq-CG-top-3 (top) and between RefSeq-ALL and RefSeq-ALL-top-3 (bottom)

#### 3.4 Tara Oceans and multi-hierarchy analysis

Comparisons with similar methods were performed in this manuscript with one index including all organism groups of interest. However, ganon provides a multi-hierarchical classification system that can use several filters interchangeably, giving more flexibility to perform analysis. This feature together with the fast build and update times allow the creation of independent indices that can be gradually updated and used in any combination.

As a use case we analyzed one sample from TARA Oceans project [5] whose organisms are known for being underrepresented in the current repositories, even with recent advances in cataloging such environment. Besides Archaea (A), Bacteria (B), Fungi (F) and Virus (V) reference sets used from the previous evaluation, we also added Plasmids (PL) and Protozoa (P) reference sets, all from RefSeq. In addition, we built an index based on a set of recently assembled contigs from Tara Oceans project (TARA) [6]. The reads from sample ERR599159 were trimmed by quality and size and one million random reads were analyzed, with the following quality trimming:
fastp=0.19.7:
length_required 95
qualified_quality_phred 20
n_base_limit 3
low_complexity_filter

We first classified the sample against the "standard" RefSeq dataset used in the previous section, achieving less than 3% of classification on the RefSeq-CG-top-3 set and less than 6% in RefSeq-ALL-top-3 (Figure 15). As expected, a higher number of reads was achieved classifying

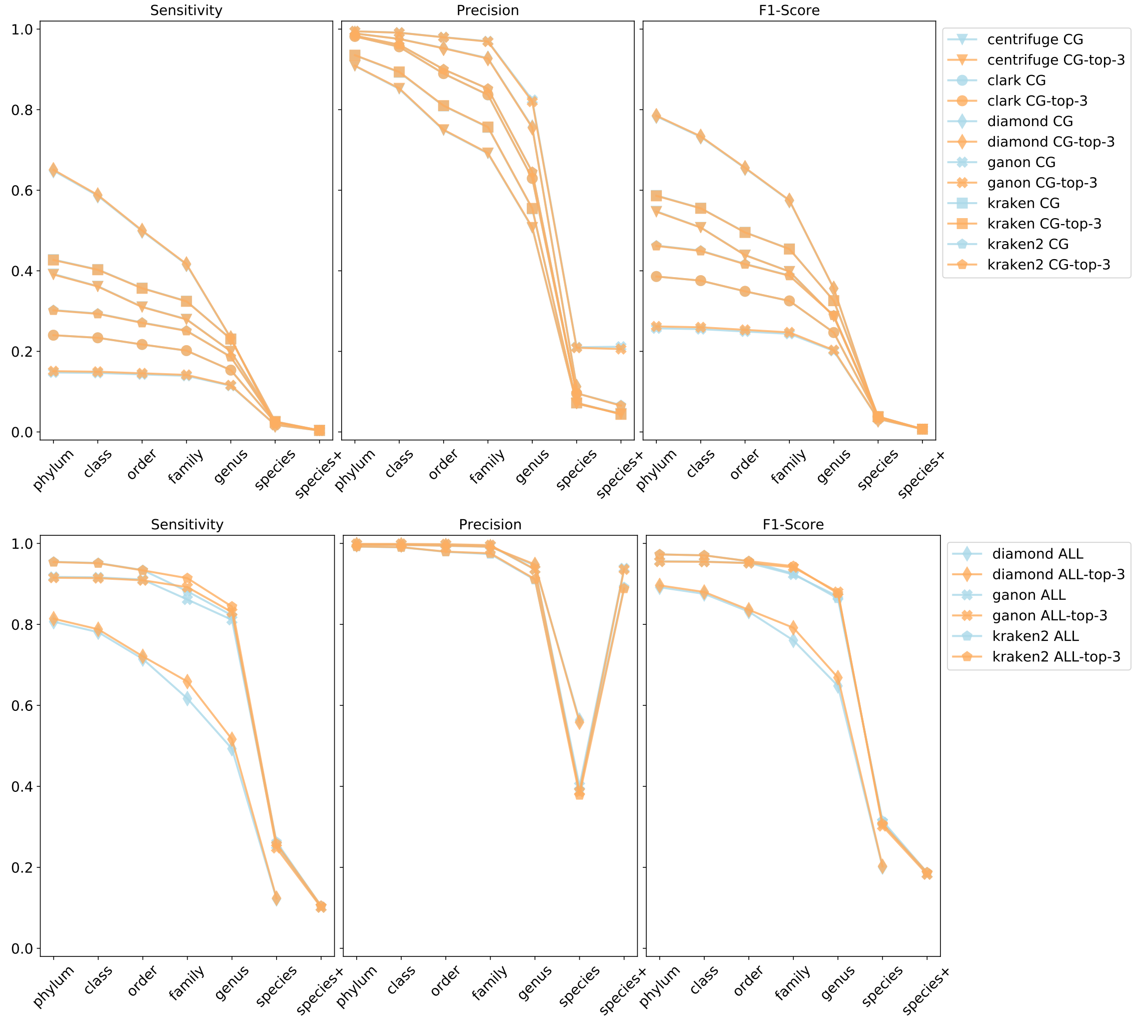

Figure 11: Rank-based precision, sensitivity and F1-Score for the real reads between complete RefSeq-CG and RefSeq-CG-top-3 (top) and between RefSeq-ALL and RefSeq-ALL-top-3 (bottom)

them against the custom TARA reference set, with slightly more than 12% of reads classified.

Since the TARA set also contains Virus and Microbes, it is hard to interpret such results independently from each other. Here we could take advantage of ganon multi-filter analysis. We ran ganon classify combining Bacteria, Archaea, Fungi and Virus as independent indices and TARA sets in one run. Such combination classified more than 15.4% of the reads. In addition we performed a multi-filter and multi-hierarchy run with ganon in the following order: first reads were classified against TARA set, second reads with no matches were classified against Protozoa and Virus. Lastly, all reads without matches were classified side-to-side against Bacteria, Archaea, Fungi and the Plasmid set. This combination achieved the highest classification rate 15.5%, mainly improving the ratio of sequences assigned to species level as show in Figure 15.

Although such analyses are possible with similar approaches, ganon's fast indexing and built-in multi-hierarchical classification allow exploratory data analysis in a very fast and convenient way. Combination of several indices in any order for classification is possible with a single command. The number of errors can be set for each index as well as individual output files, allowing diverse applications (e.g. host removal before classification). In addition. given the design of Interleaved Bloom Filters, it is possible to achieve better index to size ratio when building organism-specific indices, by clustering and adapting the size of bins and filter accordingly, speeding up the classification step.

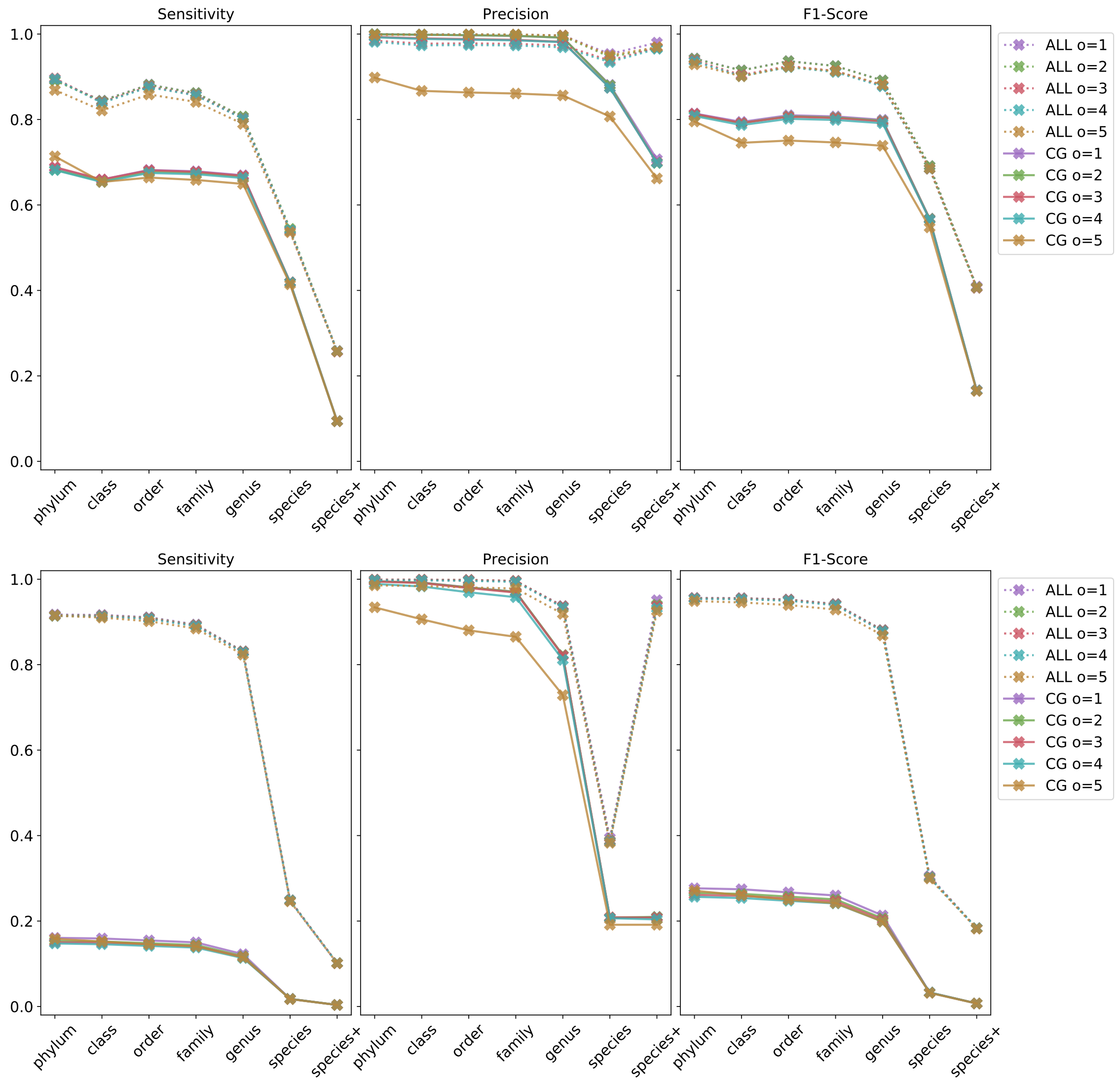

Figure 12: Variation in ganon results (rank-based) for one million random reads with different offset values ($o = 1..5$) for simulated (top) and real (bottom) reads against RefSeq-CG-top-3 (CG) and RefSeq-ALL-top-3 (ALL)

#### 3.5 Filter size

Given the design of the IBF, a trade-off between filter size and speed of classification is possible. This is done by changing the fragmentation of input sequences and reducing filter sparsity. As an example we indexed the RefSeq-CG-top-3 in three different sizes, as shown in Table 8. False positive rate (0.05), number of hash functions (3) and k-mer size (19) were kept equal for all cases. By fragmenting input sequences into more bins, ganon is able to reduce the sparsity of the filter, generating a smaller file. However, this fragmentation impacts the speed of classification. Results in terms of precision and sensitivity are unchanged as shown in Figure 16.

| filter size | # bins | classification speed (Mbp/m) |
|---|---|---|
| 32GB | 37248 | 255 |
| 48GB | 20608 | 317 |
| 64GB | 17216 | 335 |

Table 8: **Filter size and fragmentation**. RefSeq-CG-top-3 references indexed in three different sizes, showing fragmentation (# bins) and classification speed with one million random real reads (offset=3)

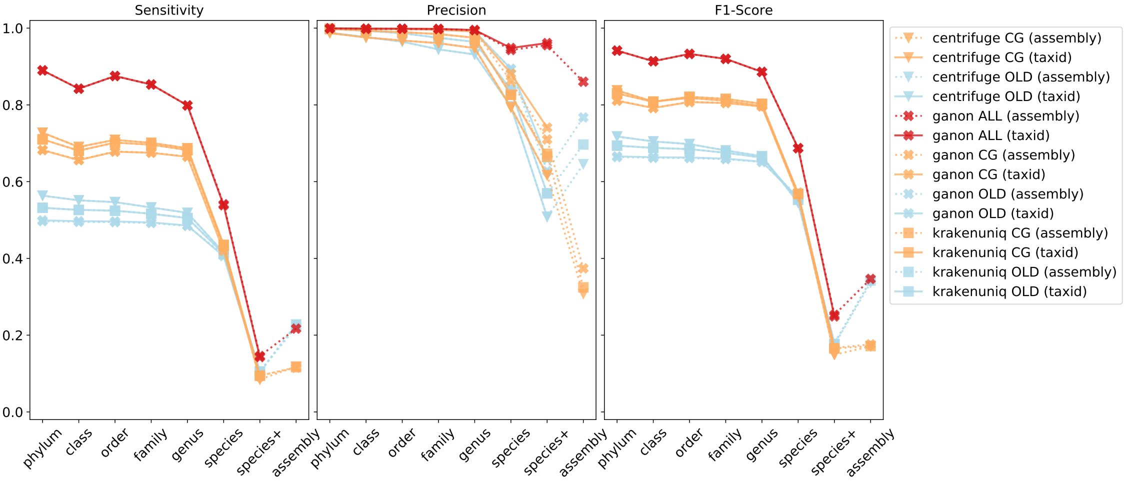

Figure 13: Rank-based precision, sensitivity and F1-Score for the simulated reads. Dotted lines show results at assembly level, while solid lines at taxonomic level for the same reference set. Kraken results at taxonomic level were used to compare to krakenuniq at assembly level (krakenuniq OLD (taxid) and krakenuniq CG (taxid)) given that they use the same base algorithm.

[5] Shinichi Sunagawa, Luis Pedro Coelho, Samuel Chaffron, Jens Roat Kultima, Karine Labadie, Guillem Salazar, Bardya Djahanschiri, Georg Zeller, Daniel R Mende, Adriana Alberti, Francisco M Cornejo-Castillo, Paul I Costea, Corinne Cruaud, F. D'Ovidio, Stefan Engelen, Isabel Ferrera, Josep M Gasol, Lionel Guidi, Falk Hildebrand, Florian Kokoszka, Cyrille Lepoivre, Gipsi Lima-Mendez, Julie Poulain, Bonnie T Poulos, Marta Royo-Llonch, Hugo Sarmento, Sara Vieira-Silva, Celine Dimier, Marc Picheral, Sarah Searson, Stefanie Kandels-Lewis, Chris Bowler, Colomban de Vargas, Gabriel Gorsky, Nigel Grimsley, Pascal Hingamp, Daniele Iudicone, Olivier Jaillon, Fabrice Not, Hiroyuki Ogata, Stephane Pesant, Sabrina Speich, Lars Stemmann, Matthew B Sullivan, Jean Weissenbach, Patrick Wincker, Eric Karsenti, Jeroen Raes, Silvia G Acinas, Peer Bork, E. Boss, Chris Bowler, M. Follows, L. Karp-Boss, U. Krzic, E. G. Reynaud, C. Sardet, M. Sieracki, and D. Velayoudon. Structure and function of the global ocean microbiome. *Science*, 348(6237):1261359–1261359, May 2015.

[6] Benjamin J Tully, Elaina D Graham, and John F Heidelberg. The reconstruction of 2,631 draft metagenome-assembled genomes from the global oceans. *Scientific Data*, 5:170203, January 2018.

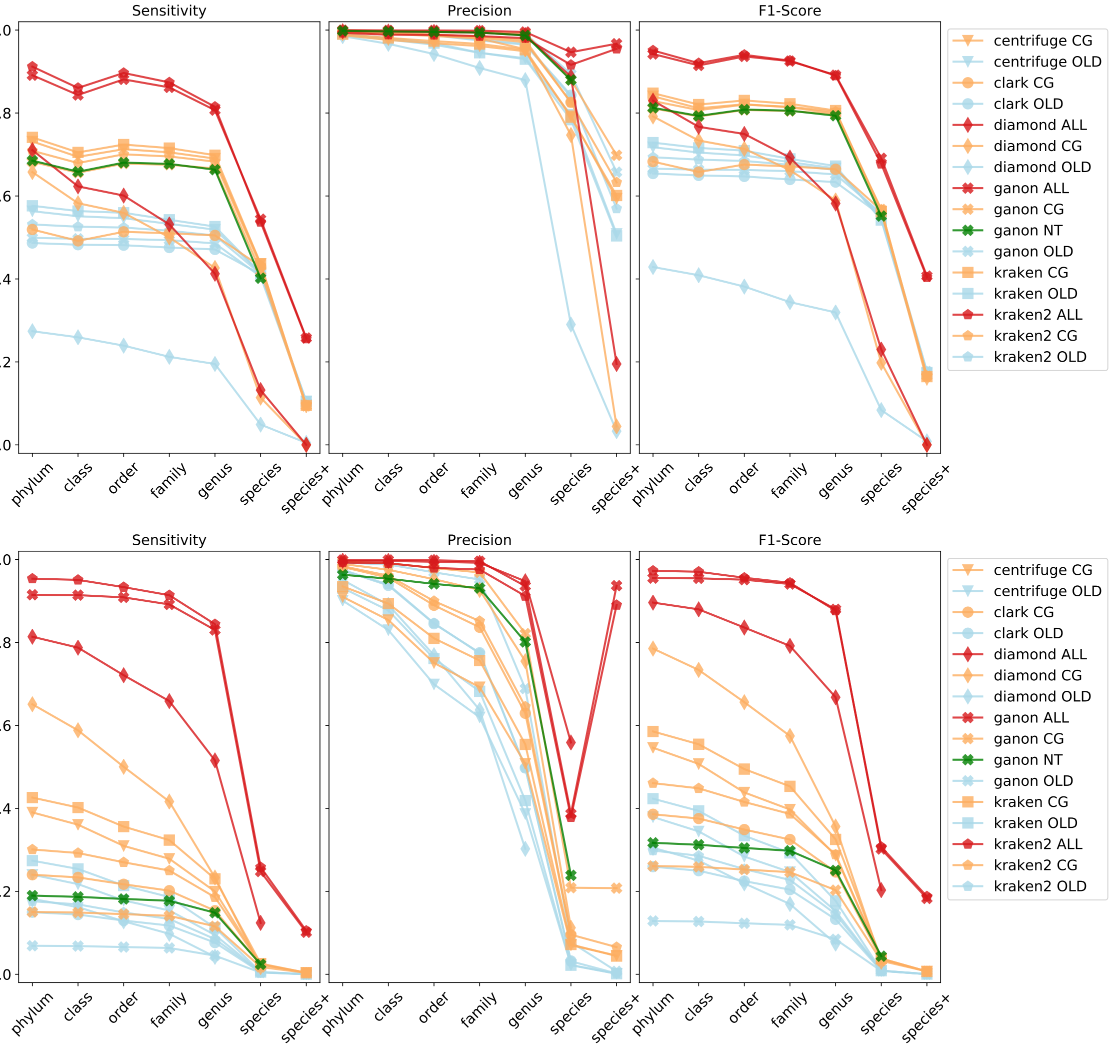

Figure 14: Rank-based precision, sensitivity and F1-Score for the a random sub-set of 1 million simulated reads (top) and 1 million real reads (bottom). In green, results against the complete NCBI-nt (not evaluated in this work) from 01-March-2019.

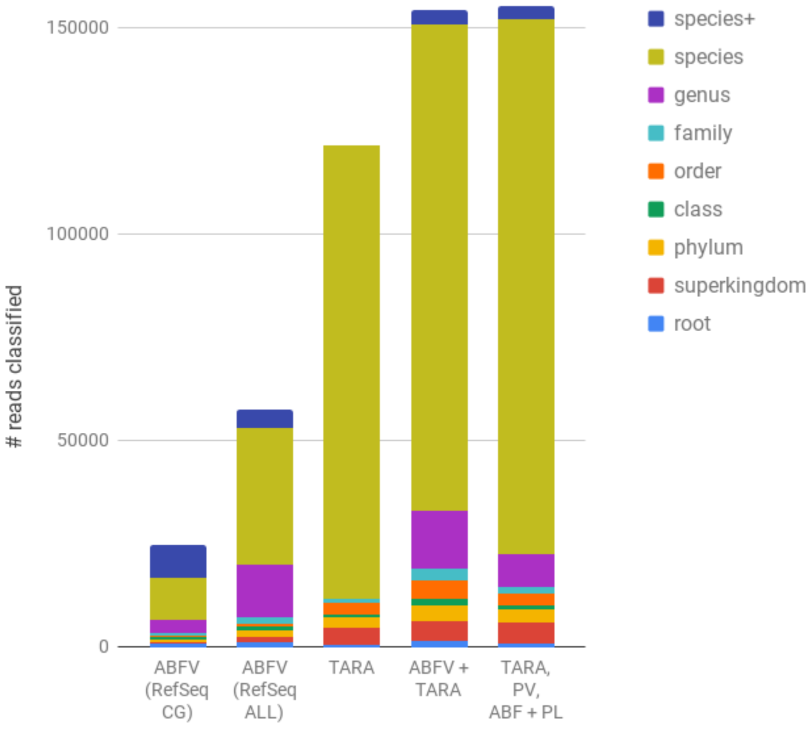

Figure 15: Tara Oceans sample analysis by taxonomic level with different combination of indices. Archaea (A), Bacteria (B), Fungi (F), Virus (V), Plasmids (PL) and Protozoa (P).

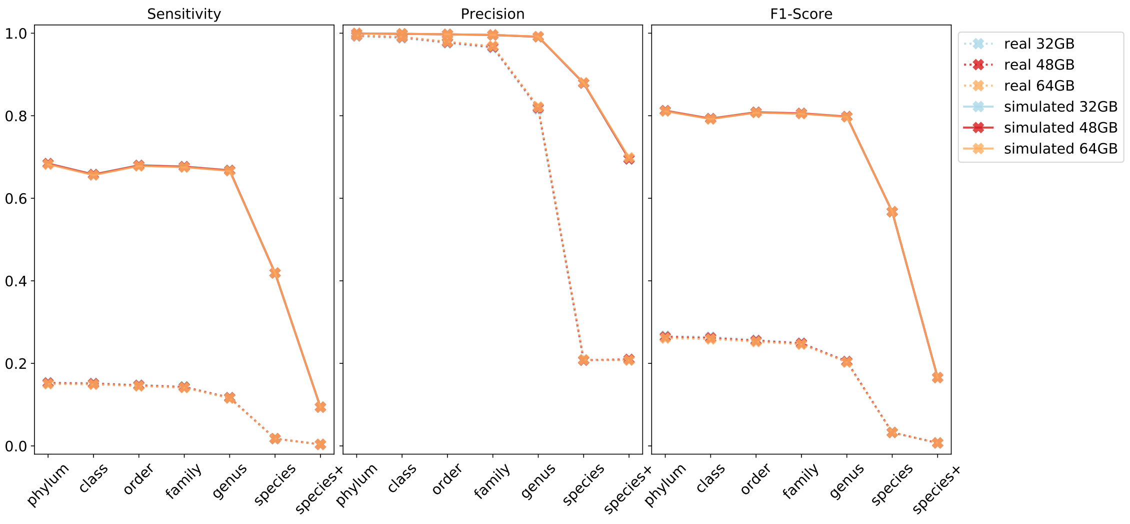

Figure 16: Rank-based metrics from the filter size variation example described in Table 8 for real and simulated reads
