## Supplementary Material 2 for "ganon: precise metagenomics classification against large and up-to-date sets of reference sequences": README.txt

The folders and files in this directory translate to names used in the manuscript as follows:

20150602_bacteria_refseq_old = RefSeq-OLD
20181219_abfv_refseq_cg = RefSeq-CG
20181219_abfv_refseq_cg_t3a = RefSeq-CG-top 3
20181219_abfv_refseq_all = RefSeq-ALL
20181219_abfv_refseq_all_t3a = RefSeq-ALL-top

cami_toy_H01 = simulated reads (taxonomic level classification)
cami_toy_H01-assembly = simulated reads (assembly level classification)
cami_challenge_H01 = real reads
