## Supplementary figures and images for "ganon: precise metagenomics classification against large and up-to-date sets of reference sequences"

### heatmap_bar.png

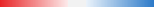
